# Chronic demyelination of rabbit lesions is attributable to failed oligodendrocyte progenitor cell repopulation

**DOI:** 10.1101/2022.01.21.477229

**Authors:** James J. M. Cooper, Jessie J. Polanco, Darpan Saraswat, Jennifer Peirick, Anna Seidl, Yi Li, Dan Ma, Fraser J. Sim

## Abstract

The failure of remyelination in the human CNS contributes to axonal injury and disease progression in multiple sclerosis (MS). In contrast to regions of chronic demyelination in the human brain, remyelination in murine models is preceded by abundant oligodendrocyte progenitor cell (OPC) repopulation, such that OPC density within regions of demyelination far exceeds that of normal white matter. As such, we hypothesized that efficient OPC repopulation was a prerequisite of successful remyelination, and that increased lesion volume may contribute to the failure of OPC repopulation in human brain. In this study, we characterized the pattern of OPC activation and proliferation following induction of lysolecithin-induced chronic demyelination in adult rabbits. The density of OPCs never exceeded that of normal white matter and oligodendrocyte density did not recover even at 6 months post-injection. Rabbit OPC recruitment in large lesions was further characterized by chronic Sox2 expression in OPCs located in the lesion core and upregulation of quiescence-associated Prrx1 mRNA at the lesion border. Surprisingly, when small rabbit lesions of equivalent size to mouse were induced, they too exhibited reduced OPC repopulation. However, small lesions were distinct from large lesions as they displayed an almost complete lack of OPC proliferation following demyelination. These differences in the response to demyelination suggest that both volume dependent and species-specific mechanisms are critical in the regulation of OPC proliferation and lesion repopulation and suggest that alternate models will be necessary to fully understand the mechanisms that contribute to failed remyelination in MS.

**Main Points:**

- Chronic demyelination in the rabbit CNS was associated with reduced OPC repopulation.
- Quiescent OPCs accumulated around the edge of rabbit lesions.
- OPC and oligodendrocyte repopulation was reduced in rabbit regardless of lesion volume.

## INTRODUCTION

Disease-modifying therapies for the treatment of relapsing-remitting multiple sclerosis (MS) are effective at limiting neuroinflammation and reduce the rate of clinical relapses (McGinley et al., 2021). However, treatment options for progressive disability in MS and agents capable of enhancing myelin repair are limited. The stimulation of remyelination has been proposed to restore neurological function and prevent neurodegeneration in regions of chronic demyelination. Although spared oligodendrocytes (OLs) may contribute to remyelination in MS and in some experimental models (Yeung et al., 2019, Duncan et al., 2018, Bacmeister et al., 2020), when large numbers of adult OLs are destroyed remyelination relies on the generation of new OLs by oligodendrocyte progenitor cells (OPCs) (Hughes et al., 2013, Zawadzka et al., 2010). The cellular processes of remyelination can be broadly divided into recruitment and differentiation phases (Franklin and Ffrench-Constant, 2017). During the recruitment phase, OPCs upregulate markers of activation, enter the cell-cycle and migrate into the demyelinated lesion resulting in tissue repopulation. The expression of activation markers, such as Sox2, are then downregulated as OPCs differentiate into OLs (Zhao et al., 2015). The differentiation phase occurs when OPC exit the cell cycle and differentiate into myelin-forming oligodendrocytes (Franklin and Ffrench-Constant, 2017). Newly generated OLs engage and ensheath denuded axons and upregulate myelin proteins resulting in the production of a thin myelin sheath with shorter internodes that is capable of restoring saltatory conduction (Lubetzki et al., 2020).

While remyelination is a common feature of early relapsing-remitting MS, remyelination failure becomes more common with increasing disease duration and age (Franklin and Ffrench-Constant, 2017, Neumann et al., 2019). Remyelination failure is thought to be due to the malfunction of one or more stages of OL regeneration: namely OPC recruitment, differentiation, or myelin formation (Franklin and Ffrench-Constant, 2017). While quiescent OPCs are associated with chronic demyelination in MS (Wolswijk, 1998), the prevailing theory in the field has been that remyelination fails in MS due to a failure of differentiation. In response, a number of drug candidates aimed at boosting endogenous remyelination through the promotion of OPC differentiation have been developed and advanced to clinical trials after successful preclinical results, notably opicinumab (anti-Lingo1) (Mi et al., 2005), clemastine (Mei et al., 2014), and bexarotene (Huang et al., 2011). Unfortunately, while all were able to improve a functional test indicative of remyelination in the optic nerve, they have largely failed to significantly improve clinical outcomes or primary imaging endpoints in initial trials (Green et al., 2017, Mellion et al., 2017, Cadavid et al., 2019, Brown et al., 2021). The paradoxical success of differentiation-promoting therapies in rodent models, but little success in clinical trials, suggests fundamental differences in the rate-limiting steps between remyelination in animal models and human MS.

A defining feature of human MS lesions is the dramatic increase in volume relative to murine models. Although highly variable in size, the median volume of an individual human MS lesions is around 100 mm^3^ (Kohler et al., 2019), whereas typical mouse lesions are < 1 mm^3^ (Jean et al., 2003, Rusielewicz et al., 2014). If the number of OLs required for successful remyelination scales linearly with lesion volume, this predicts a requirement for 100-fold more myelinating cells in MS. In the case of focal demyelination, OPCs are recruited from the surrounding white matter (Sim et al., 2002). As such, the available pool of progenitors will scale with lesion surface area which, assuming a roughly spherical lesion, increases at the square root of volume. Thus, to achieve a similar density of OPCs, a ten-fold increase in OPC generation (representing 3 or more cell divisions) and/or substantially improved migration must occur. In murine models of demyelination, the density of OPCs rapidly rises 2-3 fold above that of normal white matter shortly following lesion formation (Sim et al., 2002, Ulrich et al., 2008, Kucharova and Stallcup, 2015, Lin et al., 2006). In contrast, OPC density in MS lesions rarely exceed that of normal appearing white matter (NAWM) (Lucchinetti et al., 1999, Kuhlmann et al., 2008, Boyd et al., 2013, Moll et al., 2013, Tepavcevic et al., 2014). These observations are consistent with an inability to scale OPC recruitment as lesion size increases.

In this study, we sought to model larger volumes of demyelination which were characterized by chronic demyelination. We selected the rabbit lysolecithin model of demyelination as it was originally characterized as exhibiting chronic demyelination for at least 6 months (Blakemore, 1978, Waxman et al., 1979, Foster et al., 1980). We found that the density of OPCs never exceeded that observed in rabbit NAWM. OPC repopulation was associated with a protracted period of OPC activation and relatively poor OPC proliferation. In addition, we observed an upregulation of quiescence associated marker Prrx1 (Wang et al., 2018, Saraswat et al., 2021b) in OPCs found at the lesion border at earlier timepoints. We found that these alterations were not solely due to volume, as 10-fold smaller lesions also displayed reduced OPC density and reduced proliferation. However, small lesions were distinct in that they contained almost no proliferating OPCs. Together, these observations suggest that both volume-dependent and species-specific differences in the OPC response to demyelination contribute to the capacity of animal models to undergo spontaneous remyelination.

## MATERIALS AND METHODS

### Animals and surgery

All experiments were performed according to protocols approved by the University at Buffalo’s Institutional Animal Care and Use Committee. New Zealand White Rabbits were purchased as needed (Female, average weight 2.96 ± 0.13 kg, and average age 15.79 ± 0.53 weeks) (Charles Rivers Laboratories, Wilmington, MA). Induction anesthesia was accomplished via intramuscular injection of ketamine (35 mg/kg) and xylazine (5 mg/kg) into epaxial muscles. After loss of consciousness, supplemental heating began, and corneas were lubricated. The proximal 3 cm of the tail was shaved to provide an optimal location for the pulse oximetry probe (far preferable to use of ear or tongue). ECG leads and rectal temperature probe were placed. After pre-oxygenation via face mask, rabbits were intubated in ventral recumbency with heads and necks held in extension. A size 3.0 endotracheal tube was threaded over an 8-French urinary catheter. After placement of the catheter through the glottis, the ET tube was advanced over the catheter and into the trachea. Successful intubation was verified via immediate capnography. Upon success, isoflurane was delivered via Bain Circuit (1-3%). An IV catheter was placed (marginal ear vein), and the surgical site was shaved and prepped prior to moving the rabbit to the operating room (OR). Pre-emptive, initial analgesics were administered (0.05 mg/kg buprenorphine, subcutaneously (SC) and 1.5 mg/kg carprofen, SC).

Once moved onto the OR table, rabbits were immediately reconnected to isoflurane to ensure a suitable anesthetic plane. Then, they were transiently disconnected from the Bain Circuit to allow loose movement of the head into the stereotaxic frame (Kopf Model 902 Small Animal Stereotaxic Instrument + Kopf Model 1240 Rabbit Adaptor + Rabbit Risers, Kopf Instruments, Tujunga, CA). The endotracheal (ET) tube, was carefully moved to rest between the arms of the Kopf Rabbit Adaptor. It was important that the ET tube was not advanced beyond 12 cm to avoid bronchial obstruction. Once positioned, the capnograph and Bain Circuit were quickly reconnected. At this time further careful positioning of the rabbit’s head was possible via tooth bar, zygoma clamps, nose clamp. General anesthesia could be maintained for the remainder of the stereotaxic procedure with isoflurane. Spontaneous respirations were augmented by manual intermittent positive pressure ventilations (IPPV). 0.1 mL bupivacaine (0.25%) or lidocaine (2%, diluted 1:1 with 0.9% NaCl) was injected SC. A 7-8 cm incision was made alone the midline of the scalp to expose both bregma (intersection of the sagittal and coronal sutures) and lambda (intersection of the sagittal and lambdoid sutures). Both left and right arm Hamilton needles were filled with distilled H_2_O to prevent occlusion and ensure flow prior to lysolecithin loading. Placing the needles point on either bregma or lambda, the dorsal-ventral (DV) relationship was adjusted using the Kopf Rabbit Adaptor until bregma was 1.5 mm above lambda. The Rabbit Alignment Tool (Kopf Model 1244 Rabbit Alignment Tool) greatly aided the speed of this process, as well as ensuring minimal rotation about the principal axes (roll, pitch, yaw). The arms of the Kopf frame were adjusted to a 20° angle, and the needle points were placed on bregma. The anterior-posterior (AP), lateral-medial (LM), and DV of each arm was recorded. The following adjustments were made to these values: AP= +0.5 mm, LM= +7.7 mm, DV= −5.7 mm. Injection sites were confirmed previously via Evans Blue injection. Drill bits (Burrs for Micro Drill 19007-29, 2.9 mm, Foster City, CA) were sterilized via dry micro bead bench side sterilizer for 1 minute and cooled with room temp saline. In advance of drilling, topical articaine (4%) was dripped onto the skull and allowed to absorb and dry over drilling skull sites. Drill holes were made through the skull to the level of the meninges with constant saline stream to prevent parenchymal heat damage. A 30G needle was then used to puncture the meninges to allow for free passage of the Hamilton needle. 1% lysolecithin was then injected at a rate 0.1 μl/min, with 5-minute wait times between microliters to allow for diffusion. Once injection was completed, the needles were left in place for 20 additional minutes to allow for diffusion. Either 5 μL (large volume lesion) or 0.35 μL (small volume lesion) or lysolecithin was injected.

Post-operatively, buprenorphine was administered (0.05 mg/kg, SC) 4 hours after the initial dose, and carprofen (1.5 mg/kg, SC) was continued twice daily for a total of 4 consecutive days. Rabbits were monitored daily after surgery and assessed for: mentation, weight loss from baseline, appetite, water consumption, amount and character of feces, amount of urination, and presence of neurologic signs. Post-surgical weight loss peaked at 28 dpl, followed by a rapid recovery (data available upon request).

### Tissue processing

Animals were sacrificed at 7, 14, 21, 56, or 180 days post-lesion (dpl) by transcardial perfusion of saline followed by 4% paraformaldehyde under deep anesthesia. Following decapitation, whole brain tissue was extracted and post-fixed for 30 minutes in 4% paraformaldehyde. For cryopreservation, the tissue was first left in 1× PBS overnight, then transferred into a 7.5% sucrose solution overnight, and finally 15% sucrose overnight. Cryoprotected tissue was then frozen in optimal cutting temperature medium (Tissue-Tek). Serial, 16 μm-thick coronal sections were cut using a Leica cryostat and stored in −80 °C freezer until processing.

To identify lesion location and volume, every 10^th^ section was washed with 1× PBST (×3 by 5 min), stained with FluoroMyelin for 1 hour at room temperature (1:300, Thermo Fisher Scientific) and 4′,6-Diamidino-2-phenylindole dihydrochloride (DAPI) for 3 min (1:5000, Sigma Aldrich), and mounted in prolong gold (Thermo Fisher Scientific). Lesions were identified as regions with lesser FluoroMyelin signal and corresponding hypercellularity by DAPI, compared to surrounding white matter. Cross-sectional lesion areas were calculated for both left and right hemisphere lesions using the NIH ImageJ software, and the largest cross-sectional area for each lesion was identified as the lesion center.

### Estimation of lesion volume

Lesion volume was calculated from the measured cross-sectional areas determined by FluoroMyelin staining using Cavalieri’s estimator of morphometric volume (Rosen and Harry, 1990):

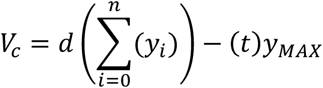

Where *V_c_* is the Cavalieri’s estimated volume, *d* is the distance between sections, *y_i_* is the cross-sectional area of section *i*, *t* is the thickness of the sections, and *y_MAX_* is the largest cross-sectional area of the lesion. Since every 10^th^ section was stained, d =160 μm, and t =16 μm.

### Immunohistochemistry (IHC)

Slides immediately adjacent to the lesion centers, identified by FluoroMyelin staining, were used for all immunohistochemical procedures. In general, sections were permeabilized for 1 hour with 1% Triton X-100 (Alfa Aesar) and 0.25% Tween 20 (Calbiochem). Sections were blocked for 1 hour with a solution containing 0.5% Triton X-100, 5% bovine serum albumin (BSA), and 5% normal donkey serum. For cytoplasmic antigens such as CC1, sections were permeabilized with a 0.1% saponin and 1% normal donkey serum for 15 minutes and blocked for 1 hour with a 5% donkey serum and 0.05% saponin. Primary antibodies utilized were goat anti-Olig2 (1:100, R&D Systems), mouse anti-CC1 (1:50, Millipore), mouse IgG2a anti-SOX2 (1:100, R&D Systems), mouse IgG1 anti-Ki67 (1:25, Fisher Scientific), rat IgG2a anti-MBP (1:300, Abcam Inc.), goat IgG Smi311 & Smi312 (1:1000, BioLegend), rabbit IgG anti-App (1:500, Thermo Fisher), rabbit IgG anti-Iba1 (1:300, Wako Chemicals), and mouse IgG1 anti-Gfap (1:400, Sigma-Aldrich). Alexa 488, 594, and 647-conjugated secondary antibodies (Invitrogen) were used at 1:500. Lesion areas were captured at 10X, 20X, and 60X magnification using an Olympus IX83 with IX3-SSU motorized stage and image tiles stitched via cellSens software. Images of immunostained sections were aligned with their corresponding FluoroMyelin stained section to accurately mark lesion boundaries, with exclusion of any portion of the lesion which extends into the gray matter. Cell counts were determined via a semi-automated, machine learning assisted, counting process (code available upon request). The Keras based convolution network was trained on a set of ~3250 CC1 positive and ~3250 CC1 negative pre-classified images (supervised learning).

### Fluorescence in situ hybridization

Fluorescence in situ hybridization was performed with probes targeting Prrx1 (GenBank accession #NM_011127.2) mRNA by using the RNAscope Fluorescent Multiplex Detection Kit (Advanced Cell Diagnostics) according to the manufacturer instructions and as described previously (Wang et al., 2018, Saraswat et al., 2021a). Fixed sections were baked at 60°C for 30 minutes, washed with ethanol, tissue pretreatment with proprietary buffers. Probes were then hybridized and fluorescent conjugated. Sections were counterstained with DAPI to visualize nuclei. Positive signals were identified as punctate dots present in nucleus and cytoplasm.

### Statistics

All statistical testing was completed using GraphPad Prism Version 8.4.0. The control group for all analyses of cell numbers and phenotypes was normal white matter (NWM) from uninjured animals, or normal appearing white matter (NAWM) from a region distant from the lesion. Groups comprised either time-points following injection of lysolecithin, regional analysis within the lesion, or small vs large volume lesions. For all reporting, an experimental unit comprised a single lesion. A minimum of 4 experimental units (lesions) per group was set *a priori*, along with a requirement for at least three animals per group. Criteria for lesion inclusion were established *a priori*. Only lesions located within internal capsule white matter were included in the analysis. No white matter lesions regardless of size were excluded from the analysis. Rabbits were allocated to each group in a randomized fashion based on availability from the vendor and surgical schedule. Experimental units for large and small lesion animals are displayed in **Table 1**. Experimental units for normal uninjured animals: 6 hemispheres, 3 animals. Total animals involved in the study, 38. Manual cell count validations for all quantitative measures were performed in a blinded fashion. Normality of data was tested in GraphPad using the D’Agostino-Pearson omnibus test. For analysis of time-series data, one-way ANOVA with Tukey’s post-hoc testing was performed. For analysis of time-series and region/volume delimited data, two-way ANOVA with Sidak’s post-hoc testing was performed. Pairwise post-hoc testing was only performed if supported by ANOVA. Only a portion of pairwise comparisons were highlighted, however full pairwise statistical comparisons are available upon request.

**Table 1.** Animal and lesion numbers per timepoint.

| Timepoint<br>(dpl) | Large Lesions |  | Small Lesions |  |
| --- | --- | --- | --- | --- |
|  | Animals | Lesions | Animals | Lesions |
| 7 | 6 | 9 | 4 | 6 |
| 14 | 5 | 8 | 4 | 4 |
| 21 | 4 | 8 | 4 | 4 |
| 56 | 5 | 7 |  |  |
| 180 | 3 | 4 |  |  |

### Data availability

The data that supports the findings of this study are available upon reasonable request.

## RESULTS

### Large lysolecithin-induced lesions in the rabbit persisted for at least 180 days

To test the hypothesis that supernumerary densities of OPCs are required for efficient remyelination we sought to characterize OPC dynamics in a model of inefficient remyelination. Lysolecithin-induced lesions in the sizable white matter tracts of the rabbit have been noted to persist for at least 180 days (6 months) (Blakemore, 1978, Foster et al., 1980, Waxman et al., 1979). To induce demyelination, we stereotactically injected lysolecithin into the periventricular white matter of adult New Zealand White rabbits (bilateral injections of 1% lysolecithin, 5 μl) (**Fig. 1A**). We first examined lesion formation at 14 days post lesion (dpl). Overlay of a mouse coronal section included for size reference. Lysolecithin induced focal demyelination at the site of injection and resulted in a clearly demarcated region of hypercellularity that precisely matched the region of myelin loss (**Fig. 1B**, white dashed line). The mean maximal cross-sectional area (MCA) and volume were greatest at early time points and significantly declined with time (volume: 14 vs 21 dpl, Tukey’s *p* = 0.032; MCA: 14 vs 56 dpl, Tukey’s *p* = 0.013) (**Fig. 1C-D**). Large regions of demyelination persisted in the rabbit even at 180 dpl. Volumetric analysis by serial section reconstruction of the lesion indicated a comet-like appearance with a relatively spherical region of demyelination corresponding to the site of injection and a tapered region extending in the posterior-ventral direction (**Fig. 1E**).

**Figure 1.**
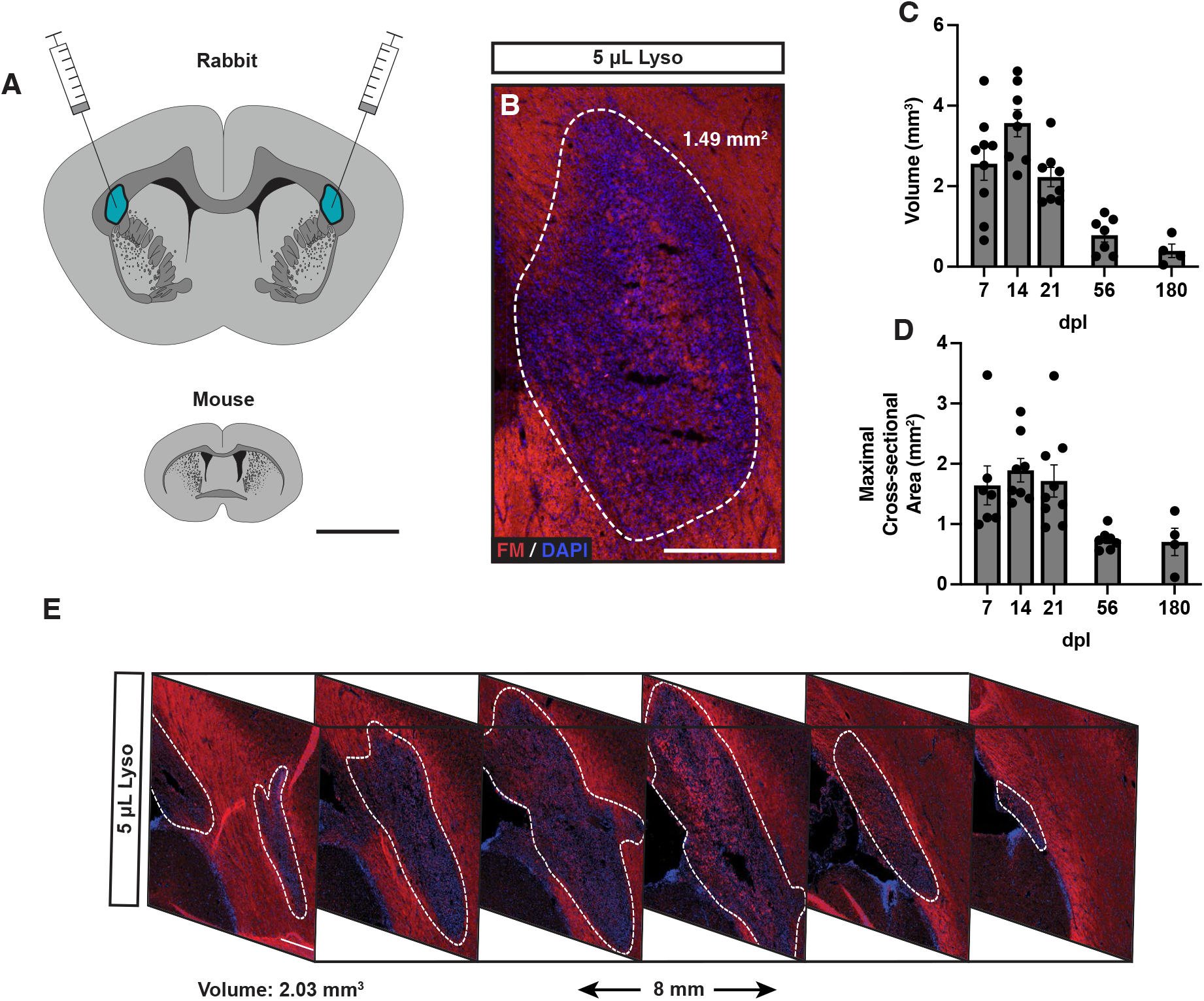
Large regions of demyelination following injection of lysolecithin into rabbit subcortical white matter persisted for at least 180 days. **A**, Scale diagram representing injection sites and white matter tract size (mouse included for reference). **B**, Lysolecithin-induced lesion at 14 days post-lesion (dpl) in the rabbit stained for FluoroMyelin (FM, red) and 4’,6’-diamidino-2-phenylindole (DAPI, blue). White dotted line indicates lesion border. **C-D**, Lesion volume (**C**) and maximal cross-sectional area (**D**) of lesions overtime. **E**, lesion reconstruction using FM and DAPI sampled every 1.6 mm, demonstrating typical lesion geometry at 7 dpl. Each point represents a single lesion. Mean ± SEM shown. Scale: 5 mm (**A**), 500 μm (**B**), 300 μm (**E**).

### Insufficient OPC repopulation in the rabbit was associated with slow and incomplete OL generation

We next sought to characterize the population dynamics of OPCs and OLs following demyelination in the rabbit brain (**Fig 2**). We used Olig2 to label the entire oligodendrocyte lineage and coexpression of CC1 and Olig2 to label post-mitotic OLs (**Fig. 2A**). Following demyelination, Olig2^+^ cell density was significantly reduced more than three-fold at 7 dpl (NWM vs 7 dpl, Tukey’s *p* < 0.0001) and never exceeded that of normal white matter (NWM) thereafter (**Fig. 2B**). The density of OPCs/immature oligodendrocytes, defined as CC1^−^Olig2^+^ cells, significantly increased between 7 and 14 dpl and remained stable at later time points (7 dpl vs 14 dpl; Tukey’s *p* = 0.009) (**Fig. 2C**). Following lysolecithin injection, the lesion was essentially devoid of OLs at 7 dpl (NWM = 877.8 ± 62 vs 7 dpl = 79.9 ± 14 cells/mm^2^) (**Fig. 2D**). By 56 dpl, OL density had partially recovered and reached a plateau but remained significantly reduced at nearly half that of NWM at 56 dpl (NWM = 877.8 ± 62 vs. 56 dpl = 453.9 ± 64.65 cells/mm^2^. Tukey’s *p* < 0.0017). The proportion of CC1^+^ OLs amongst the Olig2^+^ population has been used to infer the rate of OL differentiation. Intriguingly, while the proportion of OLs significantly increased with time until 21 dpl (7 dpl vs 21 dpl; Tukey’s *p* = 0.009) (**Fig. 2E**), the proportion of OLs did not increase at later time points suggesting that on-going OL generation may not continue after this initial period. The percentage of CC1^+^ OLs among total Olig2 remained low at 180 dpl indicating a chronic change in oligodendrocyte lineage homeostasis (NWM = 66.3 ± 3.2% vs 180 dpl = 42.2 ± 9.1%). Together, these data suggest that OPC repopulation in the rabbit lesions occurring following demyelination is insufficient to generate sufficient OLs necessary to repair these lesions.

**Figure 2.**
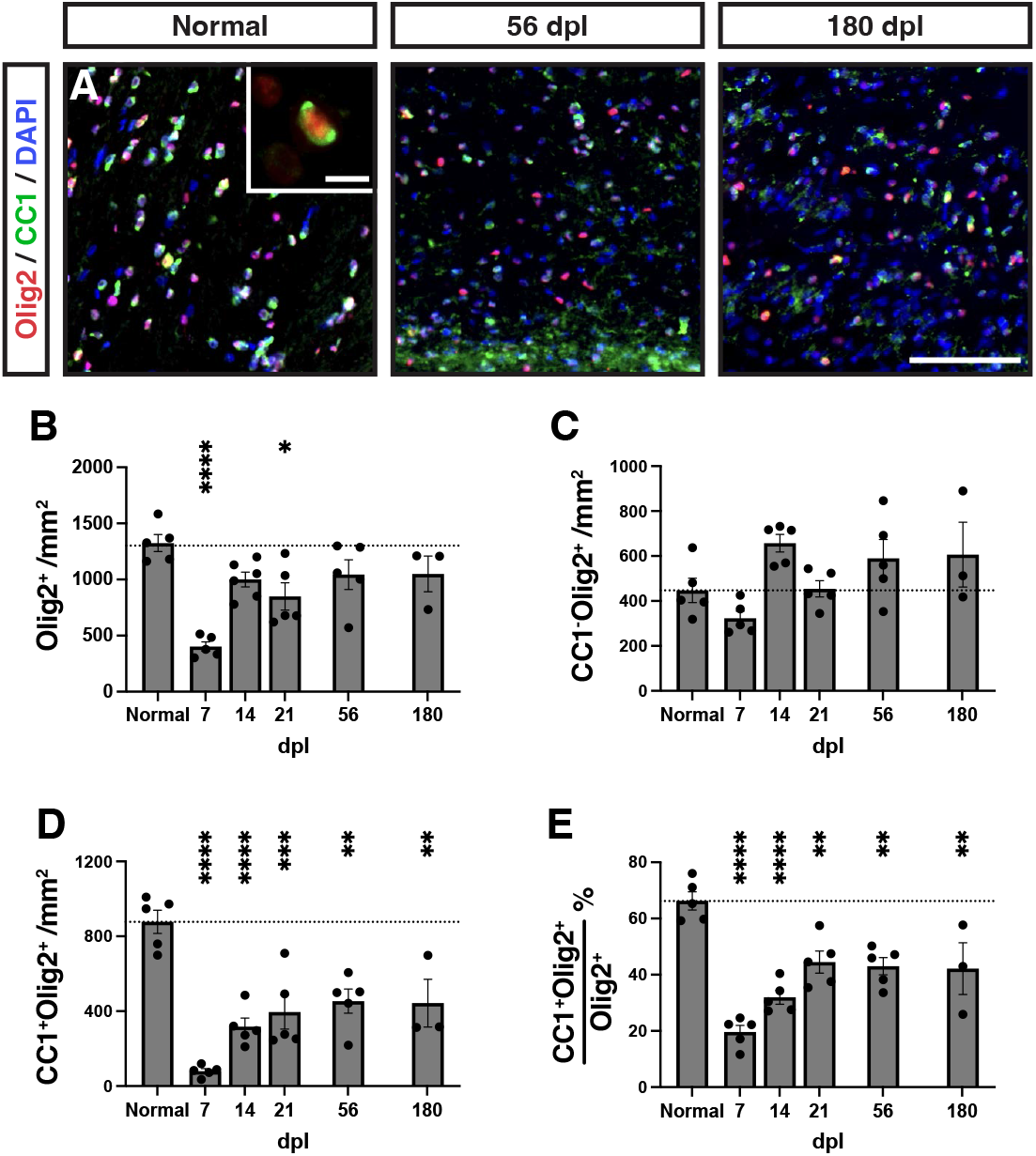
Insufficient oligodendrocyte progenitor cell repopulation and oligodendrocyte generation following demyelination in rabbit white matter. **A**, Oligodendrocyte progenitor cells (OPCs) and oligodendrocytes (OLs) were identified as Olig2^+^CC1^−^ and Olig2^+^CC1^+^, respectively. Insert shows a high magnification image of a CC1^+^ (green) OL. The density of each cell type was quantified at each time point (**B-E**). **B**, Density of Olig2^+^ oligodendrocyte lineage cells. **C**, Density of CC1^−^Olig2^+^ defined OPCs cells. **D,** Density of CC1^+^Olig2^+^ defined OLs. **E**, The percentage of CC1^+^ oligodendrocytes among total Olig2^+^ OL lineage cells. Mean ± SEM shown. Dashed line on each graph represents the mean of normal white matter (left bar, n = 6). Following one-way ANOVA, pairwise comparisons to normal were performed (Tukey’s multiple comparisons post-test). *, **, ***, and **** indicate p ≤ 0.05, 0.01, 0.001, and 0.0001, respectively. Scale: 50 μm (**A**), 10 μm (**A**, insert).

### OPC activation and proliferation were impaired following demyelination in the rabbit

Insufficient OPC recruitment in the rabbit may result from a failure of parenchymal OPCs to respond appropriately to environmental signals associated with demyelination, and could result from a disruption of OPC migration, activation, and/or proliferation. While we lacked the ability to directly measure migration, we instead assessed OPC activation and proliferation. The transcription factor Sox2 is upregulated in activated adult OPCs following demyelination and is necessary for OPC recruitment and successful remyelination (Zhao et al., 2015). As expected, in normal rabbit white matter the majority of Olig2^+^ cells were Sox2^-^ and Ki67^−^ (**Fig. 3A**). Following demyelination, recruited OPCs progressively upregulated Sox2, such that the majority of Olig2^+^cells in the lesion area coexpressed Sox2 by 14 dpl (NWM = 3.2 ± 2%, 14 dpl = 51.4 ± 3%) (**Fig. 3B**). Overall, the density of Sox2^+^Olig2^+^ OPC peaked at 14 dpl (NWM vs 14 dpl, 14 dpl vs 180 dpl, Tukey’s *p* < 0.0001 and *p* < 0.0001, respectively), and returned to control levels by 180 dpl (NWM vs 180 dpl, Tukey’s *p* > 0.05) (**Fig. 3B-E**). The pattern of Sox2^+^ expression among OL lineage cells largely corresponded with their proliferative capacity defined by Ki67 expression. The proportion of Ki67^+^ cells among the Olig2^+^ population peaked at 14 dpl and returned to baseline by 180 dpl (NWM vs 14 dpl, 14 dpl vs 180 dpl, Tukey’s *p* = 0.004 and *p* = 0.02, respectively) (**Fig. 3B-D, F**). Interestingly, the proportion of dividing Sox2^+^Olig2^+^ remained elevated until 56 dpl but never exceeded 30% (NWM vs 56 dpl, Tukey’s *p* = 0.025) (**Fig. 3G**). Together, this indicates a protracted period of OPC activation, whereby OPC proliferation fails to keep pace, and one that ultimately fails to result in supernumerary OPC repopulation.

**Figure 3.**
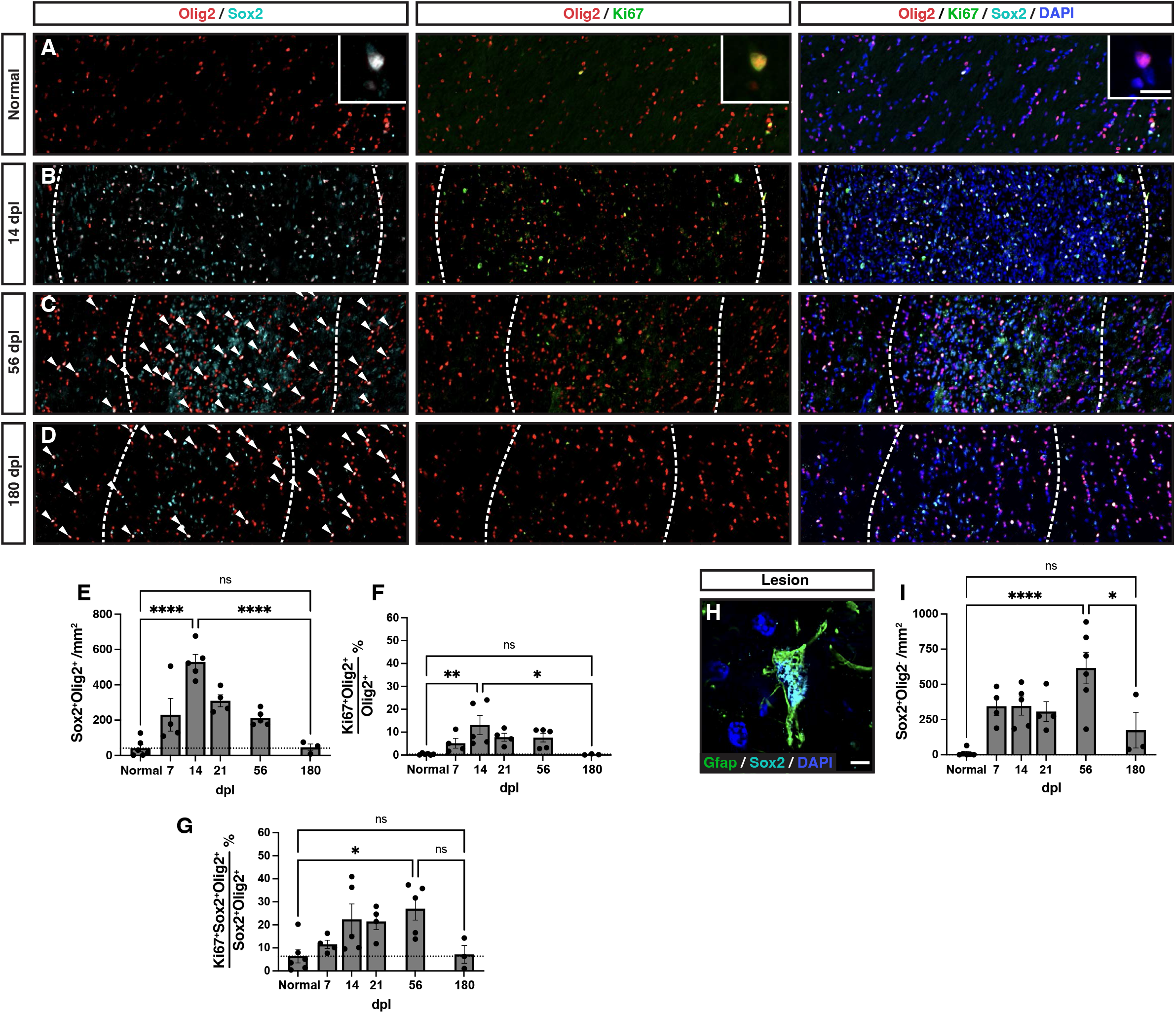
Abundant oligodendrocyte progenitor cell activation but low proliferation following injury. **A-D**, Representative images of normal white matter (**A**), or lesion at 14 days post-lesion (dpl) (**B**), 56 dpl (**C**), 180 dpl (**D**). Activated Sox2^+^ OPCs (cyan), proliferative Ki67^+^OPCs (green) were identified by colocalization with Olig2 (red). Insert shows high magnification of labelled cells. Individual Sox2^+^Olig2^+^ cells are indicated with arrows in **C** and **D**. **E**, Density of activated Sox2^+^ OPCs following demyelination. **F**, Proportion of Ki67^+^ proliferating Olig2^+^ cells. **G**, Proportion of Ki67^+^ proliferating cells among Sox2^+^Olig2^+^ activated OPCs. **H**, An example of Sox2 (cyan)-expressing Gfap^+^ (green) astrocyte within the demyelinating lesion. **I**, Quantification of Sox2^+^Olig2^−^ astrocytic cell density. Mean ± SEM shown. Dashed lines on graph represent mean of normal white matter (left bar, n =6). One-way ANOVA was performed across time points and specific pairwise comparisons are shown, between normal and peak, peak and 180 dpl, and normal and 180 dpl (other pairwise comparisons excluded for clarity) (Tukey’s multiple comparisons post-test). *, **, ***, and **** indicate p ≤ 0.05, 0.01, 0.001, and 0.0001, respectively. Scale: 200 μm (**A-D**), 10 μm (**A**, inserts), 5 μm (**H**).

### Microglial/macrophage response correlates with the pattern of OPC repopulation

Next, we examined the microglial and astrocyte response following demyelination to determine whether the failure to recruit supernumerary OPCs could be attributed to the pattern of the innate immune response. In this regard, we noted substantial upregulation of Sox2 among Olig2 negative cells following demyelination (**Fig. 3B-D**). We confirmed that these Sox2^+^ cells were largely reactive Gfap^+^ astrocytes that were concentrated within the center of the demyelinated lesion (**Fig. 3H**). A substantial number of Sox2^+^Olig2^−^ cells were observed within the lesion at all time points, with statistically greatest numbers at 56 dpl followed by a reduction at 180 dpl (NWM vs 56 dpl, 56 dpl vs 180 dpl, Tukey’s *p* = 0.045 and *p* = 0.018, respectively) (**Fig. 3I**).

At 7 dpl, both microglial (Iba1) and astrocyte (Gfap) markers were upregulated throughout the demyelinated lesion with a relatively homogenous pattern throughout the lesion area (**Fig. 4A**). At higher magnification, Iba1^+^ microglia largely adopted an ameboid morphology while astrocytes were hypertrophic and reactive in appearance (**Fig. 4A**, insets). At later timepoints both microglia/macrophage and astrocyte signals were more concentrated toward the lesion center (**Fig. 4B-D**). While Gfap immunoreactivity remained elevated through 180 days, Iba1 was more transient with a peak between 7-14 days. Consistent with this qualitative assessment, Iba1 mean florescent intensity (MFI) peaked at 7 and 14 days and was significantly elevated relative to normal appearing white matter (**Fig. 4E**) (NAWM; defined as matched uninjured corpus callosum from the same cohort of animals, n=29). Iba1 was elevated relative to NAWM at early time points up to and including 56 dpl (NAWM vs 56 dpl, Tukey’s *p* = 0.006). However, Iba1 staining was reduced at 180 dpl (7 dpl vs 180 dpl, Tukey’s *p* < 0.0001) (**Fig. 4E**). In contrast, the astrocytic response defined by Gfap area remained significantly elevated throughout the experimental time course, with significant Gfap upregulation vs. NAWM still apparent at 180 dpl (NAWM vs 180 dpl, Tukey’s *p* = 0.02) (**Fig. 4F**). Gfap upregulation was stable as no significant pairwise differences were observed between 7 dpl and any later timepoint (Tukey’s *p* > 0.05). Consistent with these observations, the total density of DAPI^+^ nuclei in the lesion area peaked at 14 dpl and declined thereafter. DAPI^+^ density was significantly decreased 56 vs. 14 dpl (Tukey’s 14 vs 56 dpl, *p* = 0.04) (**Fig. 4G**). Together, these results suggest that astroglial and microglial responses are distinct from one another and, intriguingly, that the overall microglial rather than the astroglial response corresponded to the pattern of OPC activation and proliferation.

**Figure 4.**
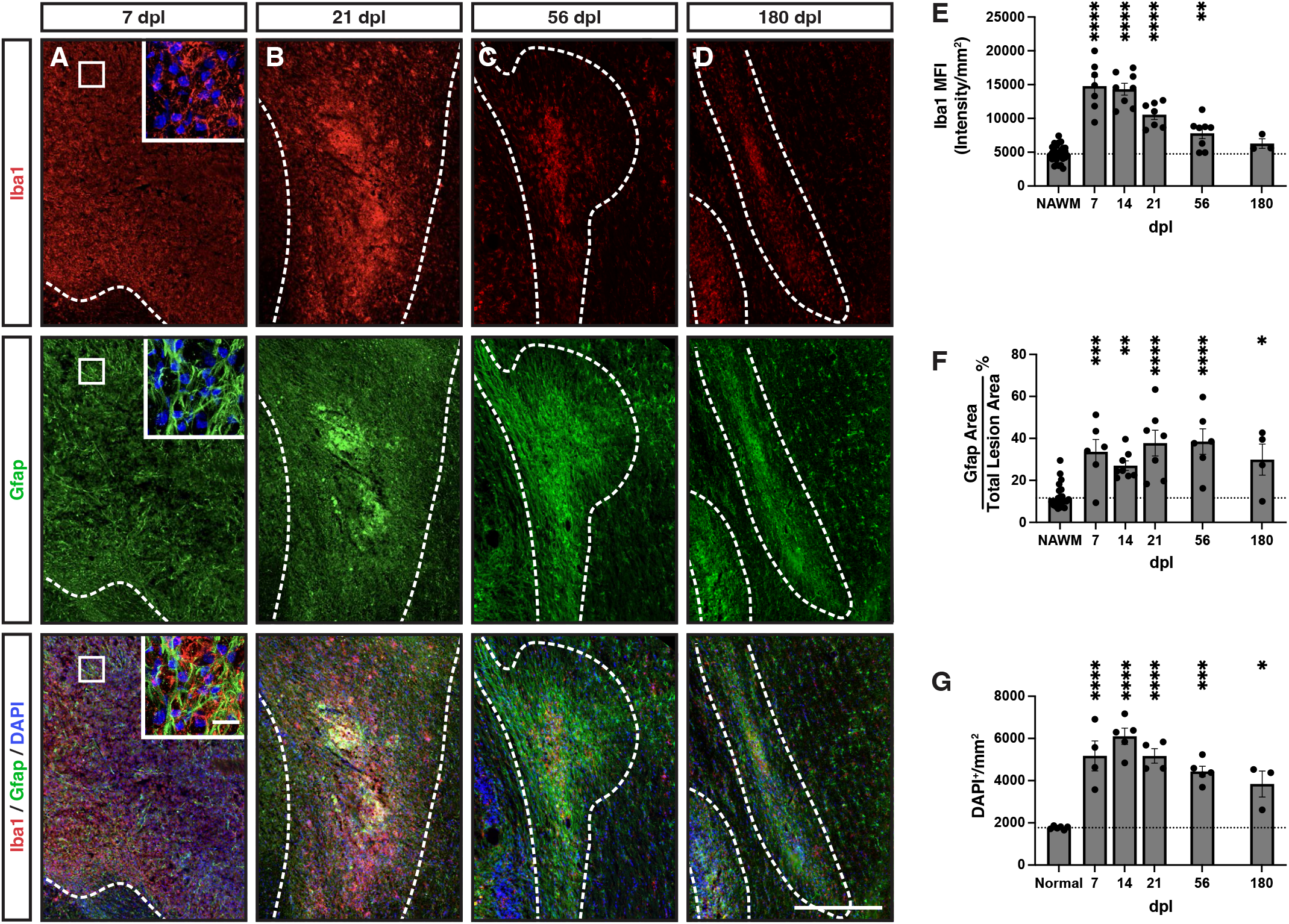
Large lesions displayed abundant gliosis, with the microglia/macrophage response greatest at early timepoints. **A-D**, Iba1 (red) and Gfap (green) were used as markers of microglia/macrophages and astrocytes, respectively, with nuclei labeled with DAPI (blue). Lesion borders (dashed line). Inserts show higher power confocal images of boxed areas. **E**, Iba1 mean fluorescent intensity (MFI). The intensity of staining in the distant corpus callosum was considered normal appearing white matter (NAWM) for quantitative comparisons and to control for batch and animal staining variability. **F**, Total Gfap^+^ area above threshold was quantified and shown as a percentage of total lesion area. **G**, DAPI^+^ cell density within the lesion (cells / mm^2^). Mean ± SEM shown. Dashed lines on graph represent mean of distant normal appearing white matter (NAWM) (n= 29) or normal uninjured animals (n=3). Following one-way ANOVA, pairwise comparisons to NAWM were performed (Tukey’s multiple comparisons post-test). *, **, ***, and **** indicate p ≤ 0.05, 0.01, 0.001, and 0.0001, respectively. Scale: 500 μm (**A-D**), 25 μm (**A**, inserts).

Interestingly, in the center of the lesion ameboid microglia/macrophages commonly formed vesicular structures which were observed at early 14 dpl (**Fig. 5A**), and later 56 dpl timepoints (**Fig. 5B**). These structures overwhelmingly represent myelin ladened microglia/macrophages, which were also present at 56 dpl, albeit at lower number (**Fig. 5C-E**). Additionally, while almost all of these vesicular Iba1^+^ structures were ladened with myelin at 14 dpl, there were a significant portion at 56 dpl which did not demonstrate immunoreactivity to myelin debris or neurofilament (**Fig. 5D-E**). As myelin debris impairs remyelination (Kotter et al., 2006), these data suggest that inefficient myelin clearance in the lesion core may influence repair in the rabbit.

**Figure 5.**
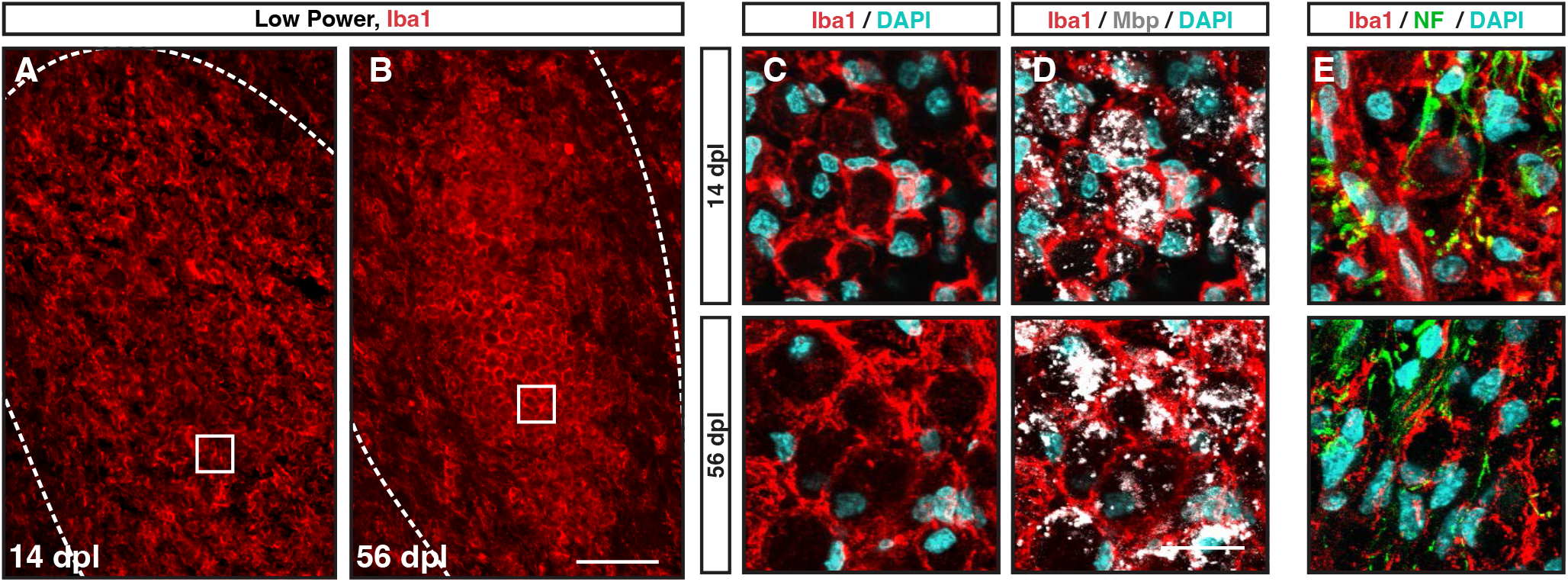
Myelin ladened microglia/macrophages were abundant within the lesion center of large rabbit lesions. **A-B**, Low power imaging revealed Iba1^+^ (red) vesicular structures filling the lesion core at 14 (**A**) and 56 dpl (**B**). **C-E**, Confocal microscopy with DAPI (blue)(**C**), myelin basic protein (Mbp, gray) (**D**, same field), and neurofilament (NF, green) (**E**) co-staining with Iba1 (red). The majority of vesicular structures surrounded by Iba1^+^ immunoreactivity contained myelin debris at 14 dpl while a small minority contained neurofilament. At 56 dpl, a substantial proportion of vesicular structures did not contain immunoreactivity to Mbp or NF. Scale: 100 μm (**A-B**), 20 μm (**C-E**).

### Large rabbit lesions display regional differences in cellularity, demyelination, and axonal loss

Toxin-induced demyelination in mouse models is characterized by a region of hypercellularity which corresponds to the extent of myelin degeneration. We compared the extent of myelin loss defined by FluoroMyelin (FM) with the relative density of DAPI^+^ nuclei between 7 and 180 dpl (**Fig. 6**). The region of demyelination, defined by loss of FM staining, was hypercellular at 7 dpl (**Fig. 6A**) and remained so thereafter. However, surrounding the region of demyelination (**Fig. 6A**, white dashed line), we also observed a region of mild hypercellularity compared to that of more distant white matter (**Fig. 6A’**, blue dashed line). We termed this region perilesion white matter (PLWM) and confirmed via quantitative analysis that the cell density was significantly elevated compared to distant NAWM (Tukey’s *p* = 0.0002) (**Fig. 6B**). We next examined the relative area of the demyelinating lesion and the associated PLWM with respect to time. Interestingly, while the area of PLWM was smaller than the demyelinated lesion at earlier timepoints, the PLWM area exceeded that of the lesion by 56 dpl and further expanded at 180 dpl (**Fig. 6C**). To investigate the potential cellular identity of cells within the PLWM, we quantified the intensity of microglial and astrocytic staining in the PLWM compared to distant NAWM. At 21 dpl, the PLWM exhibited significantly increased staining for both Iba1 and Gfap consistent with an increased number of both microglia/macrophages and astrocytes (**Supplemental Fig. 1**).

**Figure 6.**
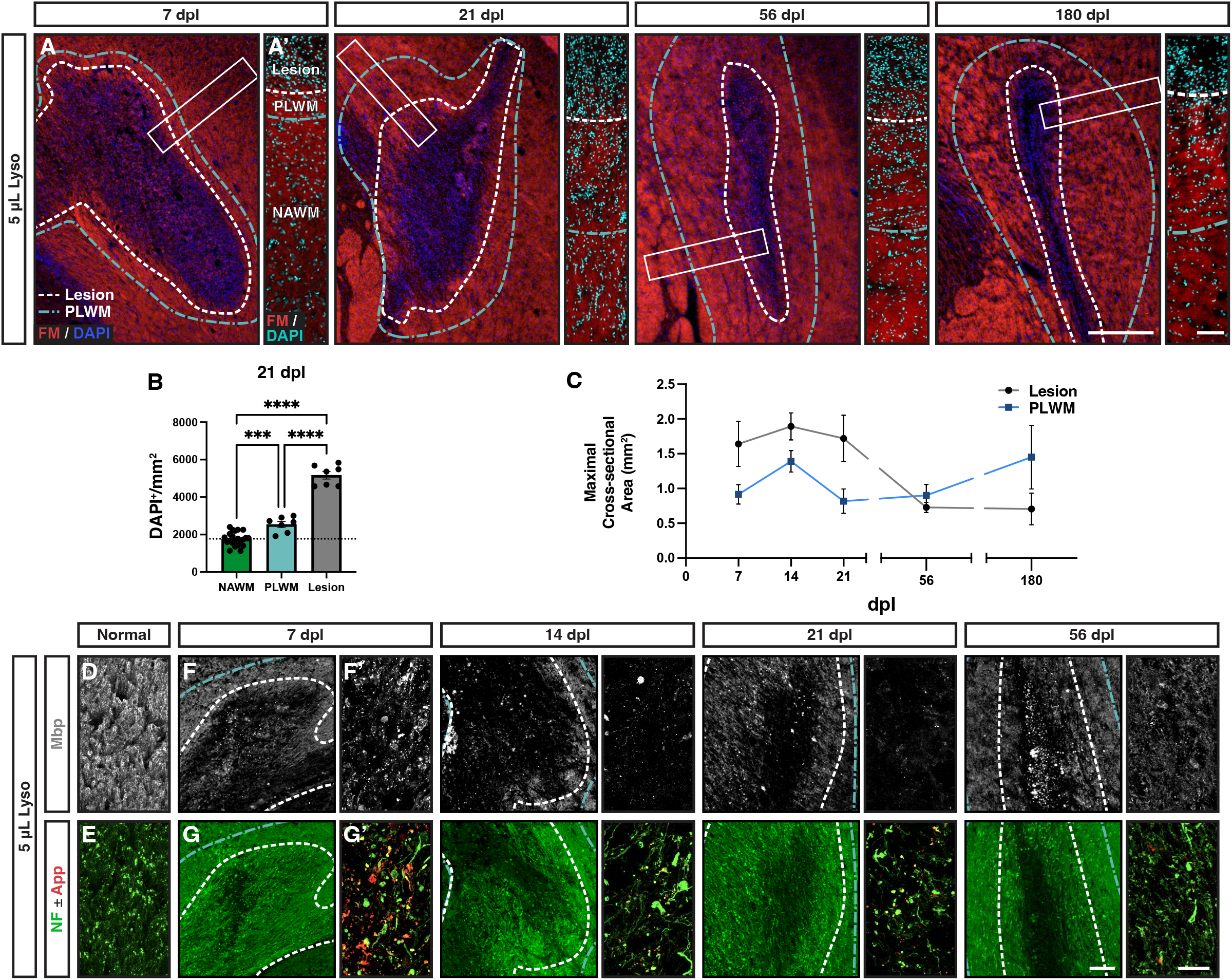
The perilesion white matter displayed hypercellularity and disturbed myelin structure. **A**, Evolution of lysolecithin-induced lesions in the rabbit over time stained for FluoroMyelin (FM, red) and DAPI (blue/cyan). White dashed line indicates lesion bounds. An area of hypercellularity extending past the lesion boundary was noted (blue dashed line), referred to as the perilesion white matter (PLWM). **A’**, Higher magnification images of the outlined areas (white rectangles) depicting the observed cellularity changes between lesion, PLWM, and more distant normal appearing white matter (NAWM). **B**, Quantification of observed changes in cellularity between the different regions at 21 dpl (n = 6-22, one-way ANOVA, F(3,38) = 147.3, *p* = 0.0001). Mean ± SEM shown. Dashed lines on graph represent mean of normal white matter. Following one-way ANOVA, pairwise comparisons to NAWM were performed (Tukey’s multiple comparisons post-test). *, **, ***, and **** indicate p ≤ 0.05, 0.01, 0.001, and 0.0001, respectively. **C**, Comparison of mean maximal cross-sectional area of lesion and PLWM areas. **D-E**, Representative confocal images of NAWM stained for myelin (Mbp, gray), and axonal markers (NF, green; App, red). **F-G**, Following demyelination at 7 to 56 dpl, lesions were imaged by widefield epifluorescence (left panels). **F’-G’**, Higher power confocal images within the lesion (right panels). Myelin loss was evident in the lesion following injection of lysolecithin and persisted thereafter. Myelin debris was present in the lesion core at all timepoints. Axonal loss was localized to the central portion of the lesion and corresponded to increased App (red) staining at 7 dpl. Scale: 500 μm (**A, F-G**), 100 μm (**A’**), 25 μm (**D-E, F’-G’**).

Next, we confirmed demyelination using myelin basic protein (Mbp) immunofluorescence that corresponded with the loss of FM staining. Compared to myelin fiber staining in NAWM (**Fig. 6D**), abundant Mbp^+^ myelin debris was observed at 7 dpl (**Fig. 6F/6F’**). The amount of myelin debris was substantially reduced by 21 dpl. By 56 dpl, there was a partial recovery of Mbp immunoreactivity within the lesion, but this was clearly distinct from NAWM (**Fig. 6F, 56 dpl**). Within the PLWM, we did not observe myelin debris at earlier timepoints (7 and 21 dpl), and the pattern and intensity of staining closely resembled NAWM. However, at 56 and 180 dpl we observed subtle changes in the PLWM with reduced Mbp intensity and uneven staining.

Consistent with primary demyelination, neurofilament (NF)-labelled axons were observed at all time points in the lesion (**Fig. 6G, Supplemental Fig. 2**). The center of the large rabbit lesion displayed a small area of localized axonal loss (9-13% of total lesion area) that remained stable in size thereafter (**Supplemental Fig. 2G**). Consistent with the loss of axons in the lesion center, we observed accumulation of App immunoreactivity specifically at 7 dpl which disappeared at later time points (**Fig. 6G’**).

### Rabbit lesions exhibit region specific gradients of OPC activation and OL differentiation

The distribution and transcriptional profile of OPCs and OLs have been shown to be dependent on their location within the lesion (e.g., core & periphery) (Boyd et al., 2013, Absinta et al., 2021). The larger rabbit lesions allowed for correlation between OPC density and the local gliotic response in specific intra-lesional regions. The lesion edge was defined as the 75 μm wide band extending into the region of demyelination from the lesion border, the remainder of the lesion will be hereafter referred to as lesion core. We examined OPC density and differentiation in each region (**Fig. 7A-E**). The lesion core and lesion edge displayed similar amounts of hypercellularity, with lesser amount of cellularity in the PLWM (**Fig. 7A**). We observed a reduction in OL density (CC1^+^Olig2^+^) that corresponded to location within the lesion, such that OL density was lowest in the lesion core and was significantly greater in the PLWM (Two-way ANOVA, region; F(2,56) = 10.72, p = 0.0001) (**Fig. 7B, 7F**). Consistent with a failure to repair regardless of region, the density of OLs in the PLWM remained significantly depressed relative to normal white matter (unpaired t test, two-tailed, NWM = 877.8 ± 62 vs PLWM at 180 dpl = 591.7 ± 110, p = 0.048).

**Figure 7.**
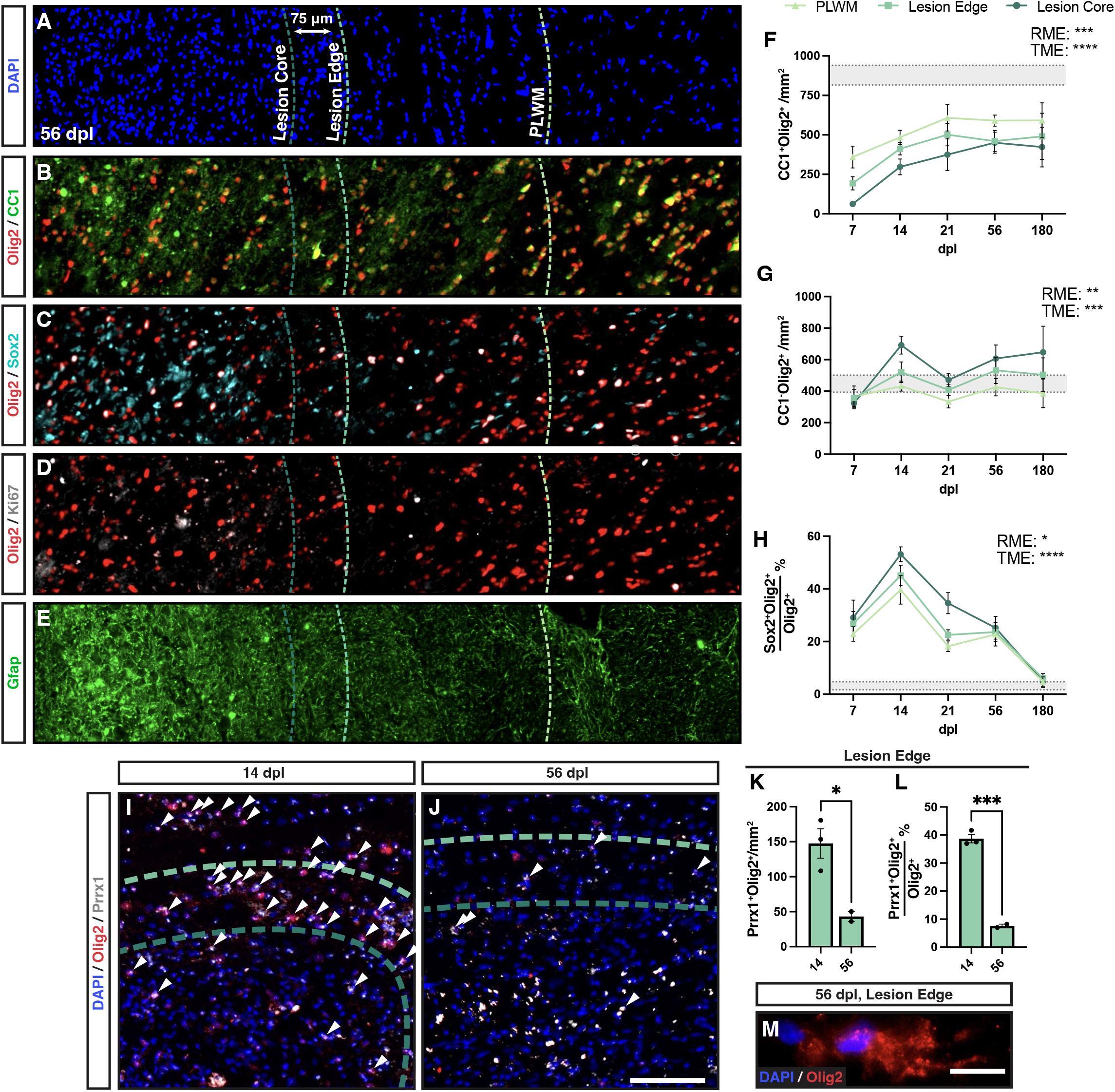
Oligodendrocyte density was reduced in the perilesion white matter, with increased expression of Prrx1 among OPCs at the lesion edge. **A-E**, Three regions were defined for regional analysis: perilesion white matter (PLWM) (light green dashed line), lesion edge (green dashed line), and lesion core (dark green dashed line). PLWM was defined as an area of mild hypercellularity and extended past the lesion border. The lesion proper was divided into the edge and core regions, with the edge defined as the outermost 75μm rim. Cross-sectional columns of a 56 dpl lesion covering the various regions stained for DAPI (**A**), OLs (Olig2^+^CC1^+^) (**B**), active oligodendrocyte progenitor cell (Olig2^+^Sox2^+^) (**C**), proliferative OPCs (Olig2^+^Ki67^+^) (**D**), and Gfap^+^ astrocytes (**E**). **F-H**, Quantification of cell density and/or proportion of total oligodendrocyte linage cells between regions for OLs (Olig2^+^CC1^+^**F**), OPCs (Olig2^+^CC1^−^, **G**), and activated OPCs (Olig2^+^Sox2^+^, **H**). Mean ± SEM shown. Horizontal bands represent mean ± 1x SEM of the normal white matter for each measurement (n = 3 rabbits). Two-way ANOVA was performed across time (TME: time main effect) and regions (RME: regional main effect). *, **, ***, and **** indicate p ≤ 0.05, 0.01, 0.001, and 0.0001, respectively. Individual pairwise comparisons not shown. **I-J**, a combination of *in situ* hybridization and immunohistochemistry was used to label Prrx1 mRNA (gray) and Olig2 protein (red). Arrowheads indicate coexpression of Olig2 and Prrx1. **K-L**, Density (**K**), and percentage (**L**) of Prrx1^+^Olig2^+^ cells in the lesion edge. Mean ± SEM shown (n=3). * and *** indicate t-test p ≤ 0.05 and 0.001, respectively. **M**, an example of a cytoplasmic Olig2 immunoreactive cell at 56 dpl. These cells were observed in most lesion sections following demyelination (~1-2 cells per section, most commonly at 56 dpl). Scale: 200 μm (**A-E**), 10 μm (**M**).

Rabbit lysolecithin lesions exhibited the greatest Olig2^+^CC1^−^ defined OPC density in the lesion core with progressively lower densities of OPCs observed in lesion edge and PLWM regions respectively (Two-way ANOVA, region; F(2,56) = 7.15, p = 0.0017) (**Fig. 7G**). The inverse pattern of OPC and OL density suggests region specific regulation of differentiation. We quantitatively examined the axonal density in lesion edge and core (**Supplemental Fig. 2H**). As expected, the lesion edge had significantly higher axonal density than core and corresponded with the inverse relationship of OPC and OL densities in these regions (Two-way ANOVA, region; F(1,30) = 21.0, p < 0.0001).

As successful remyelination in animal models is correlated with excess production of OPCs, we sought to determine why OPC density never exceeded that of NAWM in the lesion edge. Transient upregulation of Sox2^+^ by activated OPCs is required for remyelination (Zhao et al., 2015). We examined the proportion of Sox2^+^ activated Olig2^+^ cells in each region following demyelination (**Fig. 7C**). We observed the greatest proportion of Sox2^+^ OPCs in the lesion core (Two-way ANOVA, region; F(2,51) = 4.22, p = 0.02) (**Fig. 7H**). The proportion of Sox2^+^ OPCs among Olig2^+^ cells peaked at 14 dpl and declined thereafter. The relatively slow onset of Sox2 expression among rabbit OPCs suggested that other mechanisms may prevent OPC activation and proliferation at earlier time points.

### Pathological OPC quiescence may contribute to failed differentiation in the lesion edge

We have previously shown that human OPCs overexpressing the transcription factor Prrx1 enter a reversible state of quiescence that prevents their cell-cycle entry and limits their capacity to myelinate host shiverer axons (Wang et al., 2018). Similar to previous reports, following induction of demyelination Prrx1 expression was found in both Olig2^+^ oligodendrocyte lineage and other Olig2− cells (**Fig. 7I-J**). We noted a striking accumulation of Prrx1 expressing Olig2^+^ cells in the lesion edge at 14 dpl (**Fig. 7I**). This was a transient phenomenon as the density and percentage of Prrx1^+^Olig2^+^ cells decreased significantly by 56 dpl (**Fig. 7J-L**). The accumulation of Prrx1 ^+^Olig2^+^ cells at the lesion edge suggests that OPCs entering the demyelinated region rapidly upregulated Prrx1 and likely become quiescent, thereby reducing their capacity to sufficiently repopulate the lesion. Consistent with altered OPC signaling in the rabbit, we observed occasional cytoplasmic Olig2 staining in the majority of sections analyzed with cells specifically localized around the edge of the lesion (**Fig. 7M**). The appearance of cytoplasmic Olig2 has been previously attributed to cells transitioning to an astroglial fate via IFN-γ (Cassiani-Ingoni et al., 2006).

### Small volume lesions exhibited similar astrogliosis but relatively reduced microglial/macrophage responsiveness

We next asked whether the deficits in OPC proliferation and differentiation could be attributed to lesion volume alone. We created small volume lesions via injection of 0.35 μL lysolecithin. Small volume lesions were 7-fold smaller by volume than large volume rabbit lesions, and closely resembled the volume of murine lesions (Kucharova et al., 2011) (murine lesions = 0.4 mm^3^, small volume rabbit = 0.39 ±0.1 mm^3^, large volume rabbit = 3.6 ±0.3 mm^3^) (n=4 and 8, respectively) (**Fig. 8A-F, 8G**). Similar to large rabbit lesions, small volume lesions displayed equivalent levels of hypercellularity and astrogliosis (Gfap) compared to large volume lesions (**Fig. 8A-B, 8D-E**). Indeed, there were no significant differences between small and large volume lesions at any time point in terms of DAPI or GFAP (**Fig. 8H-I**). Intriguingly, we noted that small volume lesions had less intense Iba1 staining compared to large lesions (**Fig. 8C, 8F**) (Two-way ANOVA, volume: F(1,28) = 17.39, p = 0.0003). Mean Iba1 intensity was significantly decreased relative to large volume lesions at both 7 and 14 dpl (Sidak *p* = 0.0015 and 0.013, respectively) (**Fig. 8J**). Unlike larger volume lesions (1 and 5 μl), we did not observe a central region of axonal loss following injection of 0.35 μl lysolecithin (**Supplemental Fig. 2**). Axonal loss in small lesions was equivalent to that of the large lesion edge (**Supplemental Fig. 2I**).

**Figure 8.**
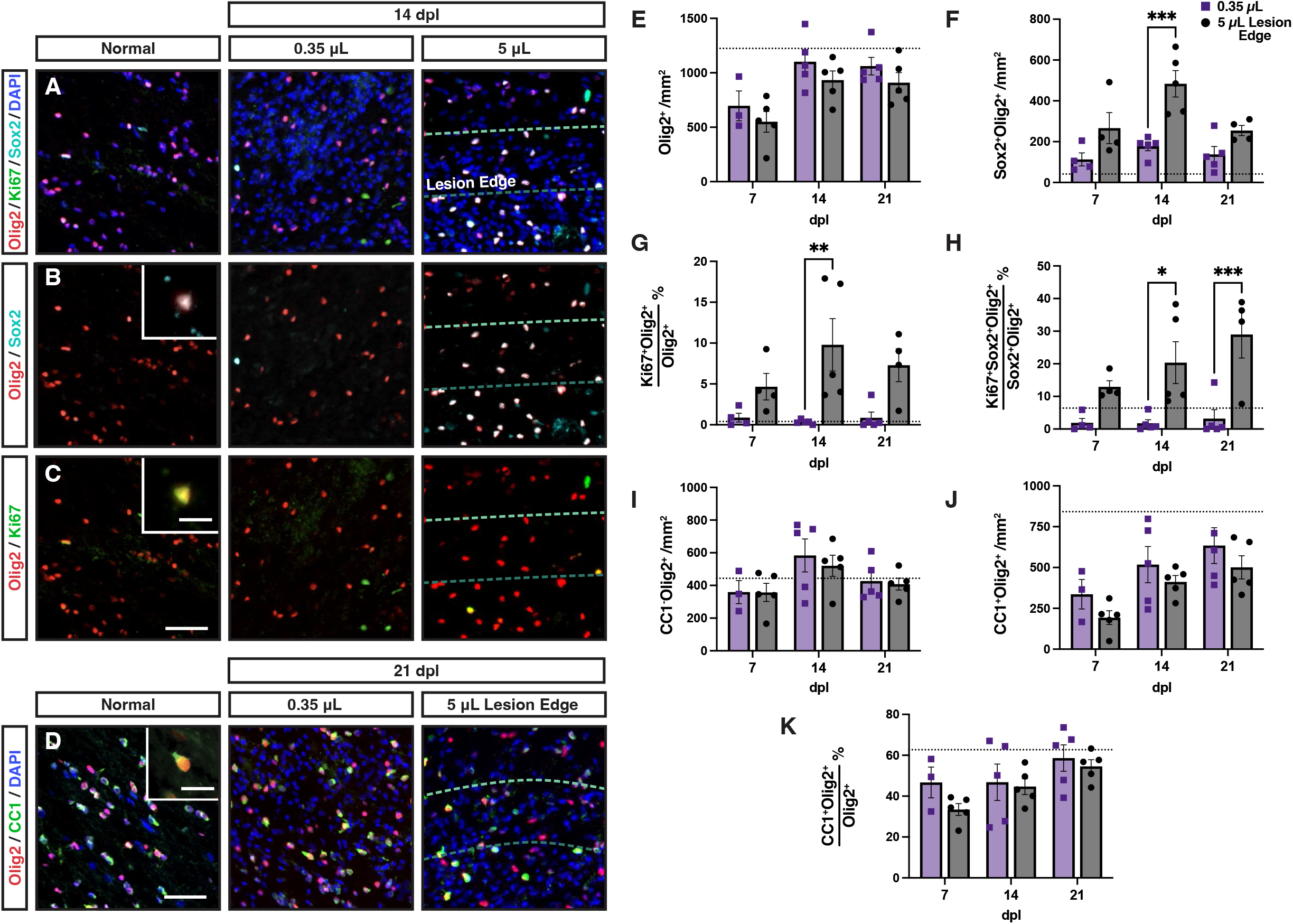
Small rabbit lesions displayed a reduced microglial response. Small (0.35 μL) (**A-C**) and large (5 μL) (**D-F**) volume lesions were created via stereotaxic injection of lysolecithin, and stained for DAPI (blue), Gfap (green), and Iba1(red). White dashed lines indicate lesion border. **G**, Lesion volume calculated by serial section reconstruction of each lesion. **H**, Lesion cell density (DAPI^+^/mm^2^). **I**, Quantification of astrocyte response. Total Gfap^+^ area above threshold was quantified and shown as a percentage of total lesion area. **J**, microglial response was quantified by mean fluorescent intensity (MFI). Mean ± SEM shown. Dashed lines on graph represent mean of distant normal appearing white matter (n= 29). 0.35 μL injections of lysolecithin produced lesions with significantly smaller volumes as compared to 5 μL at every time point (n= 4-9, two-way ANOVA, volume factor F(1,33) = 72.5, *p* < 0.0001). Small and large volume lesions displayed similar levels of hypercellularity (volume factor F(1,21) = 3.37, *p* > 0.05), and similar astrocyte responses (n= 4-9, two-way ANOVA, volume factor F(1,32) = 0.05, *p* > 0.05). However, Iba1 MFI was significantly greater in large volume lesions compared to small lesions (n= 4-9, two-way ANOVA, volume factor F(1,28) = 17.39, *p* =0.0003). Pairwise comparisons of small vs. large lesions at each time point shown for lesion volume (**G**) and microglial response (**J**) (Sidak’s multiple comparisons post-test). *, **, ***, and **** indicate p ≤ 0.05, 0.01, 0.001, and 0.0001, respectively. Scale: 200 μm.

### OPC activation and proliferation was substantially reduced in small rabbit lesions

To determine whether OPC repopulation was altered in small volume lesions, we interrogated activation and proliferation of OPCs using Sox2 and Ki67, respectively (**Fig. 9**). We compared the density of cells within the small lesion to the lesion edge of large volume lesions to exclude any effects of the central region of axonal loss. The overall density of Olig2^+^ cells following demyelination was similar between large and small lesions (Two-way ANOVA, volume; F(1,22)=3.73, *p* > 0.05) (**Fig. 9A, 9E**). However, unlike other animal models of spontaneous remyelination, Olig2 density within the demyelinated lesion never exceeded that of normal uninjured white matter regardless of lesion volume (dotted line).

**Figure 9.**
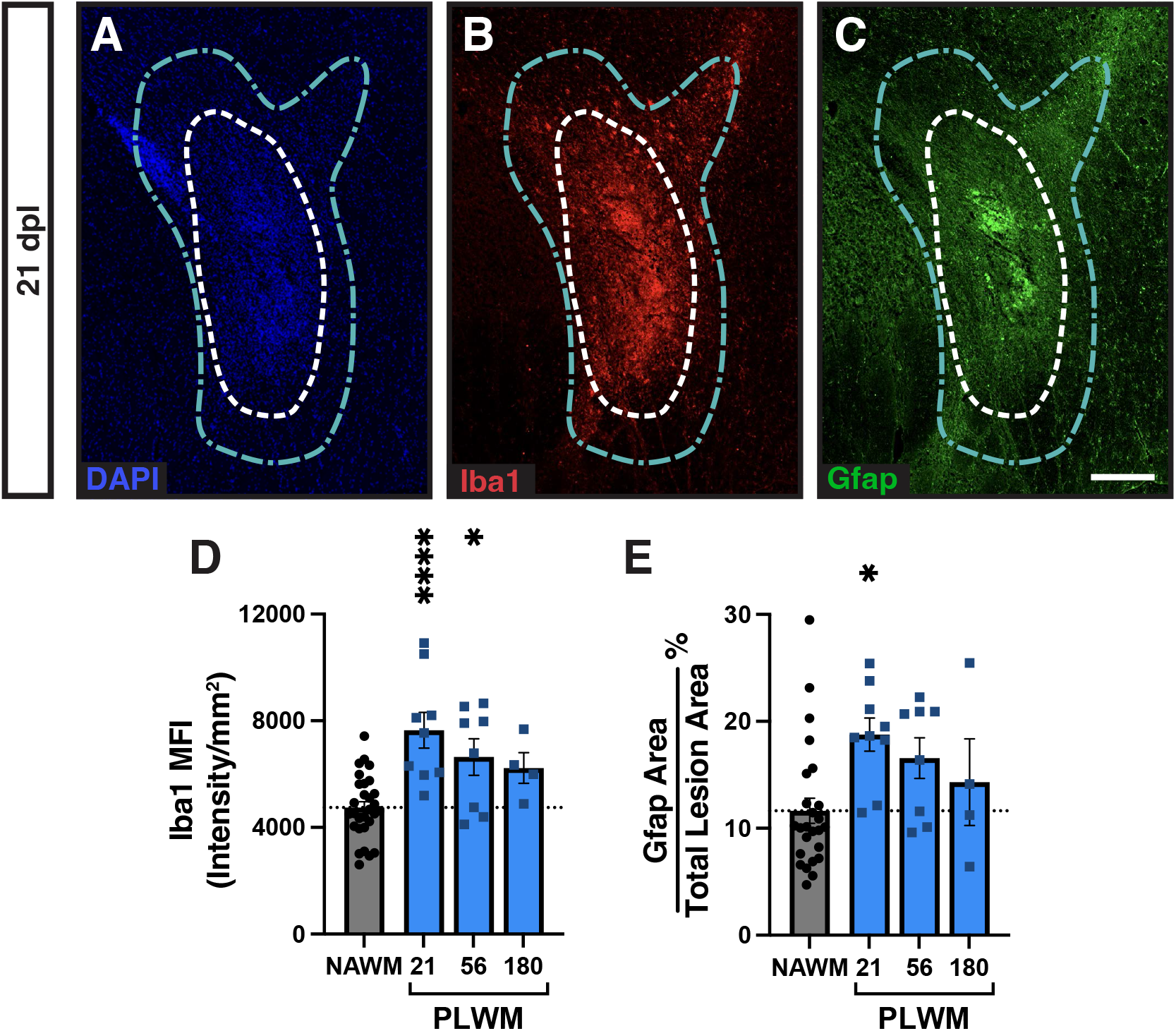
Small volume lesions displayed less oligodendrocyte progenitor cell activation and dramatically reduced proliferation. **A-C**, Activated Sox2^+^ oligodendrocyte progenitor cells (OPCs) (**B**, cyan), and proliferative Ki67^+^ OPCs (**C**, green) were identified by colocalization with Olig2 at 14 dpl in small and large volume lesions. Insert shows higher magnification of labelled cells. **D**, Oligodendrocyte (OL) differentiation was assessed by CC1 (green) and Olig2 (red) immunofluorescence. A representative CC1^+^Olig2^+^ cell is shown (**D**, inset). In large lesions, the quantification of cell density was performed in the lesion edge. **E-K**, Quantification of Olig2^+^ OL lineage cell density (cells/mm^2^, **E**), the density of Sox2^+^Olig2^+^ activated OPCs (cell/mm^2^, **F**), the percentage of Ki67-defined proliferating Olig2^+^ cells (**G**), the percentage of Ki67-defined proliferating Sox2^+^Olig2^+^ OPCs (**H**), density of CC1^−^Olig2^+^ cells (cells/mm^2^, **I**), density of CC1^+^Olig2^+^ oligodendrocytes (cells/mm^2^, **J**), and the percentage of CC1^+^Olig2^+^ oligodendrocytes among the Olig2^+^ population (**K**) in small and large lesions. Mean ± SEM shown. Dashed lines on each graph represent mean of normal uninjured white matter (n = 3 rabbits). **E-K**, Two-way ANOVA revealed a significant effects of lesion volume on activated (**F**) and proliferating (**G-H**) OPC densities (p < 0.05). The other endpoints were not significantly altered by lesion volume. **F-H**, Pairwise comparisons of small vs. large lesions at each time point were performed and shown where significant (Sidak multiple comparisons post-test). *, **, ***, and **** indicate p ≤ 0.05, 0.01, 0.001, and 0.0001, respectively. Scale: 50 μm (**A-D**), 10 μm (**B-D**, insets).

Examination of Sox2 and Ki67 revealed far more apparent volume-dependent effects (**Fig. 9B-C**). The density of Sox2^+^Olig2^+^ cells was significantly lower in small vs. large lesions following demyelination (Two-way ANOVA, volume; F(1,21) = 24.38, *p* < 0.0001). At 14 dpl, the extent of Sox2 activation was >2-fold decreased in small lesions (Sidak’s *p* = 0.0003) (**Fig. 9F**). This corresponded with a profound reduction in the proportion of actively proliferating Ki67^+^ OPCs across time points (Two-way ANOVA, volume; F(1,21) = 20.05, p = 0.0002) (**Fig. 9G**). As Olig2 expression is maintained in OLs, we next examined the fraction of Ki67^+^ cells among Sox2^+^Olig2^+^activated OPCs (**Fig. 9H**). Very few, <1%, of these OPCs expressed Ki67 suggesting that OPC proliferation is highly dependent on lesion volume in the rabbit and suggests that OPC repopulation in small rabbit lesions is principally due to migration from NAWM.

Although proliferation was substantially lower in small volume lesions, there was surprisingly no significant difference in the density of CC1^−^Olig2^+^ OPCs between small and large volume lesions (Two-way ANOVA, volume; F(1,22) = 0.25, p > 0.05) (**Fig. 9I**). Large lesions with 7-fold greater volume requires approximately 2-3 more cell divisions to achieve the same density of OPCs. The generation of CC1^+^ OLs was not significantly different between small lesions and the edge of large lesions (CC1 ^+^Olig2^+^ cell density; Two-way ANOVA, volume; F(1,22) = 3.45, p > 0.05) (**Fig. 9D, 9J**). Likewise, the rate of OL generation was equivalent between small and large lesions (Two-way ANOVA, volume; F(1,22) = 1.81, p > 0.05) (**Fig. 9K**). These results suggest that OPC activation and proliferation is highly influenced by lesion volume in the rabbit. Together, these contribute to a failure to coordinate OPC recruitment with differentiation and together lead to chronic demyelination.

## DISCUSSION

Unlike murine models, spontaneous remyelination is inefficient in rabbit models with chronic demyelination persisting at least 6 months following injection of lysolecithin (Blakemore, 1978, Foster et al., 1980), and following induction of experimental autoimmune encephalomyelitis (EAE) (Williams et al., 1982, Prineas et al., 1969). Although reported over 40 years ago, the mechanisms underlying the failure of rabbit remyelination have not been investigated. Here, we show that the OPC response to demyelination is fundamentally altered in the rabbit brain relative to typically employed rodent models. In most rodent models, OPC density far exceeds that of normal white matter during the recruitment phase of remyelination. In contrast, rabbit OPC density slowly recovers following demyelination but never exceeds that observed in normal white matter. Deficient OPC proliferation and subsequently poor OPC repopulation in large rabbit lesions was associated with an upregulation of quiescence-associated genes and this corresponded with delayed and insufficient oligodendrocyte (OL) generation. In contrast, small rabbit lesions were essentially devoid of dividing OPCs and did not efficiently upregulate Sox2 in response to demyelination. As OPC repopulation was reduced in rabbit small lesions compared to mouse lesions of equivalent size, our results also highlight the potential importance of species-dependent differences in the response to demyelination (Dietz et al., 2016).

Parenchymal adult OPCs retain the capacity to generate new OLs and are largely considered to be the primary source of newly generated OLs following demyelination in the adult CNS (Serwanski et al., 2018, Zawadzka et al., 2010). In mouse models of toxin-induced demyelination and EAE, the OPC recruitment phase of remyelination is not typically rate limiting (Franklin and Goldman, 2015). Furthermore, approaches to improve OPC recruitment have not meaningfully altered the rate of myelin regeneration in focal mouse lesions. For example, overexpression of PDGF-AA drives enhanced OPC recruitment following demyelination and yet this had no discernable effect on the rate of remyelination (Woodruff et al., 2004). The progenitor pool in small rodent models is especially capable of efficient recruitment and occurs following systemic depletion either by X-irradiation (Chari and Blakemore, 2002) or pharmacogenetic depletion (Xing et al., 2021) and, importantly, after demyelination. Repeated focal demyelination is not sufficient to deplete the progenitor pool (Penderis et al., 2003) and only sustained depletion of OL lineage cells using 12-week treatment can prevent efficient remyelination in the cuprizone model (Mason et al., 2004). The relative success of OPC recruitment in the mouse is exemplified by the rapid regeneration of OPCs following demyelination, with a 2-3 fold increase in density over normal white matter (Sim et al., 2002, Ulrich et al., 2008, Kucharova and Stallcup, 2015, Lin et al., 2006). In contrast, rabbit lesions regardless of size are characterized by a failure of OPC repopulation such that the density of OPCs never exceeds that of normal tissue and the characteristic over-population of OPCs that occurs in the mouse and rat is absent. The rate of OPC repopulation is also comparatively delayed, reaching peak density only after 14 days. Whereas mouse and rats reach peak progenitor density within the first week (Sim et al., 2002, Fancy et al., 2004). While the dynamics of OPC recruitment in humans is difficult to ascertain from postmortem tissue, MS lesions rarely display OPC densities that exceed that of NAWM (Lucchinetti et al., 1999, Kuhlmann et al., 2008, Boyd et al., 2013, Moll et al., 2013, Tepavcevic et al., 2014). Demyelination of the optic nerve in non-human primates similarly lacks the abundant recruitment of OPCs that is observed in murine models, with densities of OPCs and OLs not recovering until 9 months after injection of lysolecithin (Sarrazin et al., 2022). Fascinatingly, even at 9 months when OLs and OPC densities have recovered, few axons show evidence of remyelination (Sarrazin et al., 2022). Together these data suggest that OPC recruitment in the rabbit model more closely resembles that observed in MS and non-human primates.

While our data identify a substantial deficit in the ability of rabbit OPCs to repopulate regions of demyelination, we also noted a relative failure in the ability of OPCs to undergo differentiation. We observed a substantial decrease in the proportion of CC1^+^ OLs among the entire Olig2^+^ lineage relative to mouse/rat model systems. In the lesion core, we observed fewer OLs compared to other regions, and this corresponded with increased expression of markers of tissue injury (including OPC, microglial, and astrocytic activation and axonal loss). Together, this suggests the presence of a proinflammatory environment in the lesion core that contribute to failed OL differentiation. An alternative hypothesis is that the failure of recruitment itself contributes to failed differentiation. *In vitro* OL differentiation is dependent on the density of OPCs (Rosenberg et al., 2008) and, similarly, following transplantation human OL differentiation is correlated with the local density of human OPCs (Dietz et al., 2016). This suggests that below a certain threshold local density of OPCs, the process of OL differentiation itself may be limited. Consistent with this hypothesis, remyelinated lesions and active lesions in MS, which have a higher propensity for remyelination, tend to have higher densities of OPCs than chronic demyelinated lesions (Boyd et al., 2013). Lastly, at least within the central region of axonal loss, the failure of OL differentiation may be due to the absence of an appropriate axonal substrate. This might also account for the low density of OLs and relatively high density of OPCs found in the lesion core of large rabbit lesions. Likewise, lysolecithin-induced lesions of non-human primate optic nerve induced widespread axonal loss similar to the central region of axonal loss in the rabbit and display similarly low densities of OLs (Sarrazin et al., 2022). However, axonal loss is unlikely to explain the reduced density of OLs observed in edge of large rabbit lesions or across small volume lesions where axonal loss is not as apparent. Chronic demyelination is observed following injection of 2.5 – 5 μL lysolecithin into the rabbit spinal cord at 6 months (Blakemore, 1978). While this is likely recapitulated in this study (brain, 5 μL), it is possible that remyelination is more efficient in small (0.35 μL) lesions or those localized to the cerebral white matter. It is worth noting that the density of OLs in small lesions was not significantly greater than the lesion edge of large lesions and reduced myelin content assessed by fluoromyelin remained evident at 21 dpl suggesting a similar outcome. However, future ultrastructural analyses will be needed to elucidate this.

Several pieces of evidence in the current study point toward the importance of species differences in OPC recruitment. Contrary to our initial hypothesis, the recovery of OPC density in small volume lesions (0.4 mm^3^) was not improved relative to large volume (4.0 mm^3^) lesions and the peak density of OPCs was unaffected by lesion volume. This suggests that lesion volume is not the principal determinant of OPC recruitment. In contrast, OPC autonomous differences were apparent between the rabbit and mouse. Sox2 is upregulated by OPCs following demyelination in the mouse brain and is necessary for precursor proliferation (Zhao et al., 2015). On the other hand, downregulation of Sox2 is required for subsequent differentiation. In murine models, Sox2 is rapidly upregulated in the majority of OPCs within a week of lesion formation. In contrast, OPCs in the rabbit are slow to upregulate Sox2, failing to peak for at least 14 days. Sox2 expression persists for several weeks in the rabbit but is not associated with continued OPC proliferation suggesting a pathologic state of OPC activation. Furthermore, the transcriptional regulator Prrx1, a transcription factor associated with OPC quiescence (Wang et al., 2018), was specifically upregulated in numerous rabbit OPCs located at the lesion border. Prrx1 induces a reversible state of quiescence inhibiting proliferation and thereby preventing efficient myelination (Wang et al., 2018). Prrx1 itself is regulated by both interferon-γ and BMP signaling (Wang et al., 2018, Saraswat et al., 2021b) and increased Prrx1 expression in the rabbit suggests that the lesion environment may differ substantially from the mouse brain. Intriguingly, we also observed prominent examples of cells expressing cytoplasmic Olig2 expression in the rabbit lesion that is associated with an astroglial switch in stem/progenitor cells (Setoguchi, 2004) and observed in rare cells in EAE (Cassiani-Ingoni et al., 2006). This is not observed in mouse lysolecithin lesions and further supports species-specific differences in signaling. OPC autonomous differences between species have been suggested by several other studies. For example, adult mouse OPCs are more than twice as migratory compared to adult human OPCs (Bribian et al., 2020). Following transplantation, human OPCs take 8-12 weeks to generate myelin-forming OLs (Sim et al., 2011, Windrem et al., 2004, Windrem et al., 2008, Buchet et al., 2011), while equivalent rodent cells typically complete the process of differentiation in 3-4 weeks (Baron-Van Evercooren and Blakemore, 2004). Lastly, when human OPCs are transplanted into mouse brain, the human cells outcompete their mouse counterparts suggestive of substantial differences in cell biology and signaling between species (Windrem et al., 2014). Together, these results support the hypothesis that species differences are critical determinants of the OPC response to demyelination.

Our analysis of small and large rabbit lesions revealed interesting differences in the progenitor response to demyelination. The edge of large lesions and small lesions demonstrated similar densities of OPC and OLs at every timepoint studied. Unlike large lesions, Ki67^+^ OPCs were rarely observed in small lesions suggesting that the rate of proliferation was substantially less and strongly influenced by the amount of tissue injury. In the current study, we could not directly measure the contribution of migration to the general failure of OPC recruitment. However, as we observed a similar density of cells in small and large lesions but with the absence of proliferation in small lesions, this implies that OPC repopulation in small lesions may have been largely dependent on migration from surrounding tissue rather than local expansion. This corresponded with a much lower level of Sox2-expression amongst OPCs in small lesions and suggests an altered transcriptional state dependent on lesion volume. In addition to volume-dependent differences in the progenitor response, we also observed differences within individual lesion such that significantly higher densities of OPCs were observed in the lesion core. Proliferating Sox2^+^Olig2^+^ cells were also largely found in the lesion core and this corresponded with enhanced Iba1 and Gfap staining suggesting that the gliotic core was in part driving the sustained OPC proliferation observed in that region. In contrast, we found increased generation of OLs at the lesion border, a pattern commonly found in MS (Hess et al., 2020). The potential mediators of these differences are outside the scope of the current study but further support the importance of the local environment in the determination of OPC state and its capacity for both proliferation and differentiation.

In summary, the rabbit model provides evidence for both species and volume dependent effects on the cellular mechanisms of remyelination. Unlike mouse/rat models in which OPC and oligodendrocyte densities far exceed normal white matter densities following demyelination, in the rabbit these cells repopulate the lesion at a slower rate and never reach supernumerary densities. When comparing mouse and rabbit lesions of equivalent size, we found that activation of Sox2 occurs more slowly, and that proliferation is almost absent in the rabbit. These processes are themselves volume dependent as prolonged OPC proliferation and activation occurs in large volume rabbit lesions. The failure of OPC proliferation, and subsequently OPC recruitment, in the rabbit recapitulates the extent of OPC recruitment commonly observed in the majority of MS lesions, as well as non-human primates, and thereby provides a suitable test bed for therapeutic approaches aimed at improving the migration and proliferation of OPCs. As such, we propose that the simplicity and accessibility of the rabbit model along with the clear differences in lesion environment and patterns of OPC recruitment and differentiation will provide a vital complementary approach for preclinical testing of remyelination therapeutics and aid in the successful development of clinical interventions aimed at promoting myelin regeneration.

## Acknowledgements

This work was supported by the Department of Defense Congressionally Directed Medical Research Programs (W81XWH-21-1-0387), NINDS grant (R01NS104021), National Multiple Sclerosis Society (RG-1701-26750 and PP-1706-28080), the Kalec Multiple Sclerosis Foundation, and the Change MS Foundation. JJP received additional support from NIGMS (R25GM09545902), NCATS (UL1TR001412-S1), and New York State Department of Health (Empire State Stem Cell Fund) through the Stem Cells in Regenerative Medicine Fellowship (NYSTEM C30290GG). We acknowledge support provided by the Center for Computational Research at the University at Buffalo for use of the academic compute cluster for training the Keras-based convolution network. We also thank JA17 for is his support and motivation during this project.

## Author Contributions

Conception and design of the study: FJS and JJMC. Acquisition and analysis of data: all authors. Drafting the manuscript or figures: FJS and JJMC. Study supervision: FJS.

---

**Supplemental Figure 1.**
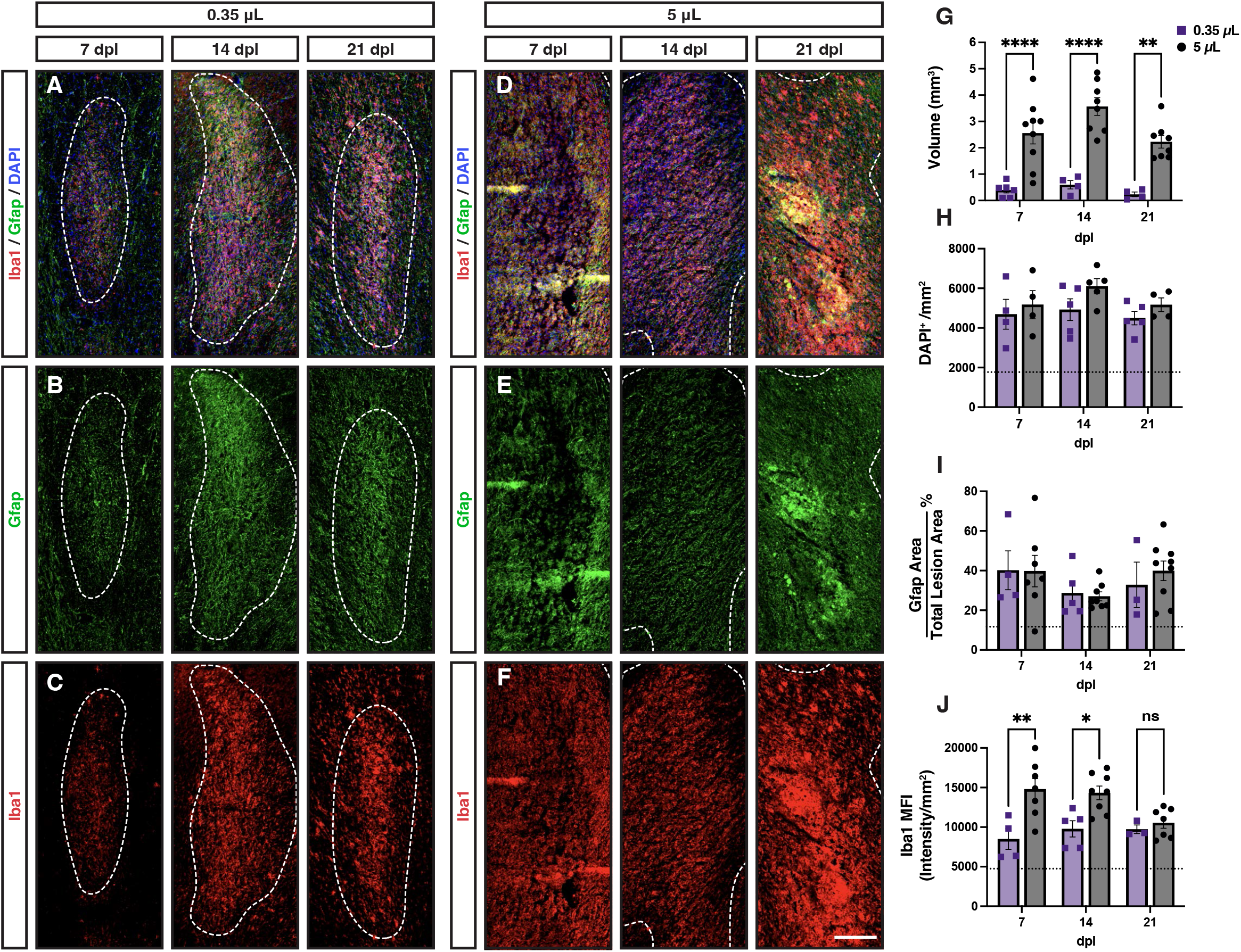
Markers of the innate immune response were increased in the perilesion white matter adjacent to demyelination. The white matter directly adjacent to the lesion area, termed the perilesion white matter (PLWM), and defined by increased cellularity compared to normal white matter exhibited increased staining for markers of microglia and astrocytes following demyelination. White dashed line indicates lesion bounds, and the PLWM was identified as an area of DAPI hypercellularity (**A**, blue) extending past the lesion boundary (blue dashed line). Iba1 was used to identify microglia (**B**, red) and astrocytes identified using Gfap (**C**, green). **D-E**, Quantification of Iba1 mean fluorescence intensity (MFI) (**D**), and Gfap % area (**E**). Mean ± SEM shown. Dashed lines on each graph represent mean of normal appearing white matter (NAWM). One-way ANOVA revealed a significant effect of time on both Iba1 and Gfap. Pairwise comparisons to NAWM were performed at each dpl (Tukey’s multiple comparisons post-test). *, and **** indicate p ≤ 0.05, and 0.0001, respectively. Scale, 250 μm.

**Supplemental Figure 2.**
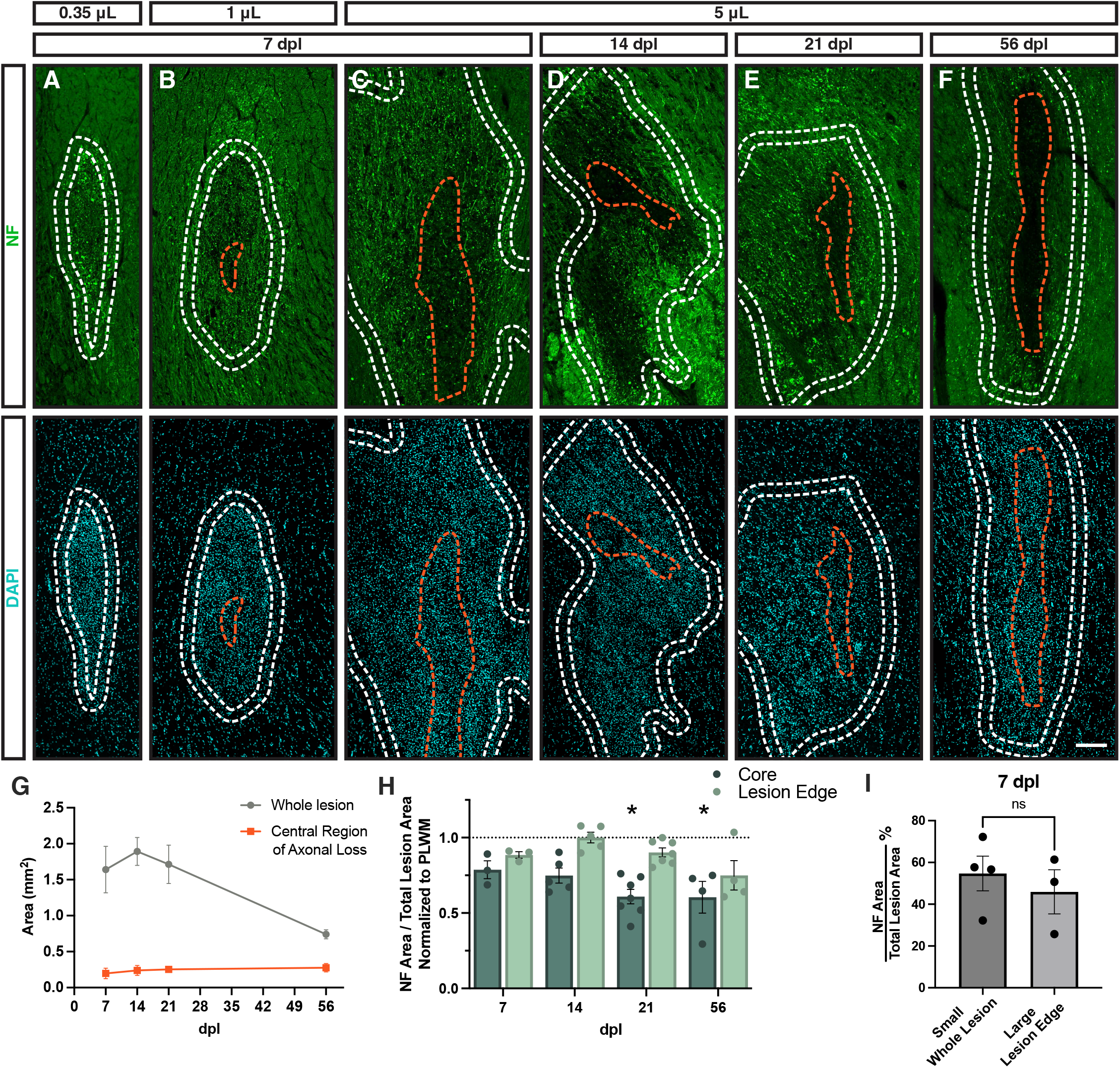
Axon deficient center was dependent on lysolecithin injection volume. Axonal density following demyelination was assessed by neurofilament immunofluorescence (green) as a function of volume of lysolecithin injected and days post lesion (dpl). DAPI (blue) hypercellularity was used to define the lesion border (outer white line). Neurofilament-expressing axonal fibers were observed throughout the lesion following injection of 0.35 μL (**A**) at 7 days post-lesion (dpl). With injections of 1 μL (**B**) and 5 μL (**C**), a central region of axonal loss was observed at 7 dpl (orange lines). The area of central axonal loss increased with injection volume. **C-F**, the central area of axonal loss persisted until 56 dpl. The area of the central region of axonal loss (orange line) compared to the total lesion area (grey line) (**G**). The size of the central region of axonal loss remained stable with time, while the overall lesion area reduced. **H**, Axonal density in the lesion edge was compared to the lesion core (which contained the central region of axonal loss). Quantitative estimation of axonal density following 5 μL lysolecithin injection was performed using a thresholding approach. Mean ± SEM shown. Dashed lines on each graph represent mean of normal appearing white matter (NAWM). Two-way ANOVA showed a significantly greater axonal density in the lesion edge than core (n = 8-14, region main effect p < 0.0001) and indicated a time dependent effect on axonal density (time main effect p = 0.014). Pairwise comparisons of 14, 21 and 56 dpl with 7 dpl identified a significant decrease in axonal density at 21 and 56 dpl in the lesion core only (* indicates Sidak p ≤ 0.05). **I**, the axonal density in small lesions was directly compared to the axonal density in large lesion edge at 7 dpl. Mean ± SEM shown. There was no significant difference (t-test p > 0.05). Scale, 200 μm.

**Supplemental Figure 3.**
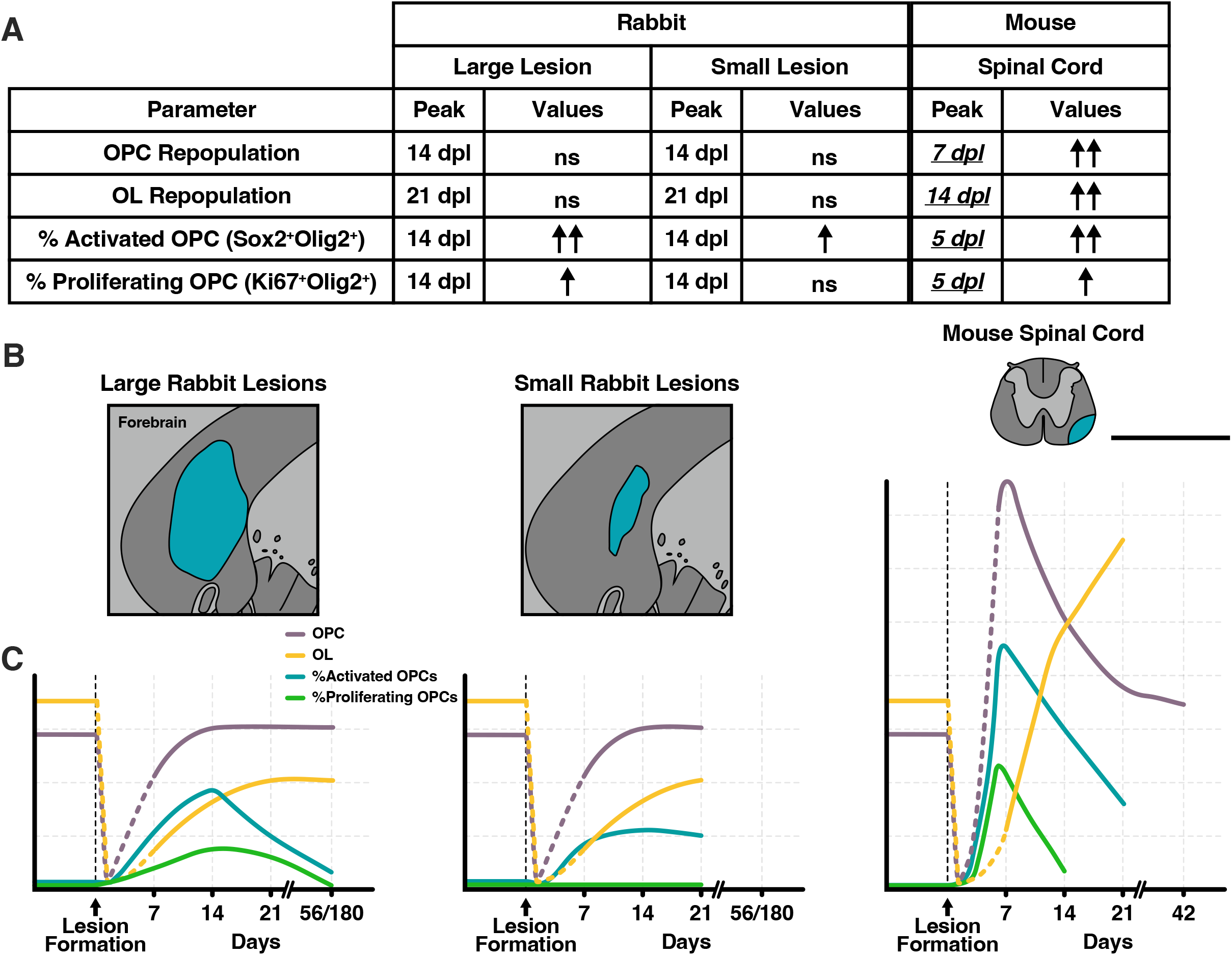
Summary of oligodendroglia population dynamics in small and large lesions, compared to mouse. Meta-analysis of rabbit large and small lesions alongside equivalently sized mouse spinal cord lesions. Comparative mouse data obtained from previously published quantification of oligodendrocyte progenitor cell (OPC)/NG2 density (Garay et al., 2011, Kucharova and Stallcup, 2015), oligodendrocyte (OL)/Plp1 (Fancy et al., 2009), Sox2^+^Olig2^+^OPCs and Ki67^+^Olig2^+^ OPCs (Zhao et al., 2015 and unpublished data). **A**, Table summarizing the effects of species and lesion volume on various cellular parameters following demyelination. The time of peak density in days post lesion (dpl) as well as the density relative to normal uninjured white matter are presented for OPC repopulation, OL repopulation, activated OPC density (Sox2^+^), and proliferating OPC density (Ki67^+^). ↑ and ↑↑ indicate significant increases in density relative to uninjured normal white matter. ns = non-significant from normal white matter density. **B**, Scale diagrams comparing large and small rabbit lesions and mouse spinal cord lesions. **C**, Profiles of overall OPC density and the relative density of Sox2^+^ activated and Ki67^+^proliferating OPCs. Scale, 2 mm.

